# Engineering PEG10 assembled endogenous virus-like particles with genetically encoded neoantigen peptides for cancer vaccination

**DOI:** 10.1101/2024.04.25.591213

**Authors:** Ruijing Tang, Luobin Guo, Tingyu Wei, Tingting Chen, Huan Yang, Honghao Ye, Fangzhou Lin, Yongyi Zeng, Haijun Yu, Zhixiong Cai, Xiaolong Liu

## Abstract

Tumor neoantigen peptide vaccines hold potential for boosting cancer immunotherapy, yet efficiently co-delivering peptides and adjuvants to antigen-presenting cells in vivo remains challenging. Virus-like particle (VLP), which is a kind of multiprotein structure organized as virus, can deliver therapeutic substances into cells and stimulate immune response. However, the weak targeted delivery of VLP in vivo and its susceptibility to neutralization by antibodies hinder their clinical applications. Here, we firstly designed a novel protein carrier using the mammalian-derived capsid protein PEG10, which can self-assemble into endogenous VLP (eVLP) with high protein loading and transfection efficiency. Then, an engineered tumor vaccine, named ePAC, was developed by packaging genetically encoded neoantigen into eVLP with further modification of CpG-ODN on its surface to serve as an adjuvant and targeting unit to dendritic cells (DCs). Significantly, ePAC can efficiently target and transport neoantigens to DCs, and promote DCs maturation to induce neoantigen-specific T cells. Moreover, in mouse orthotopic liver cancer and humanized mouse tumor models, ePAC combined with anti-TIM-3 exhibited remarkable antitumor efficacy. Overall, these results support that ePAC could be safely utilized as cancer vaccines for antitumor therapy, showing significant potential for clinical translation.

## Introduction

Neoantigens, originating from non-synonymous mutations, incomplete splicing, translation of alternatives, or posttranslational modifications, are highly immunogenic because they are absent in normal tissues and can evade central thymic tolerance mechanisms(Schumacher et al., 2019). Hence tumor neoantigen vaccines have been demonstrated as a promising and safe strategy in the treatment of various solid tumors (Ott et al., 2017; Cai et al., 2021; Chen et al., 2022). Typically, neoantigen peptides mixed with adjuvants are formulated into tumor neoantigen vaccines and directly administered to patients. Upon uptake of these antigen peptides and adjuvants, dendritic cells (DCs) are stimulated to activate neoantigen-specific CD8^+^ T cells, which in turn eliminate tumor cells and impede tumor progression (Li et al., 2017; Ott et al., 2017). However, the free antigen peptides and adjuvant inside the body following systemic administration tend to experience the rapid clearance and enzymatic degradation that would significantly limit their antitumor efficacy.

Lentiviruses offer an excellent approach to cross cell membranes by interacting with specific host cell receptors through viral envelope glycoproteins (Sundquist et al., 2012). To ensure safe gene delivery without permanent integration into the human genome, the protein of interest (POI) is always directly fused to lentiviral structural proteins for transduction (Voelkel et al., 2010; Kaczmarczyk et al., 2011; Cai et al., 2014). Specifically, the POI can be inserted into the C-terminal of matrix protein (MA), core capsid (CA), nucleocapsid (NC), or even integrase (IN) with a protease site, allowing for independent functions upon cleavage by viral protease (Voelkel et al., 2010). For example, in the delivery of CRISPR-Cas9 ribonucleoproteins (RNP), the lentiviral gag domain, including MA and NC, were fused with Cas9 and mixed with the full-length gag-pol domain for virus-like particle (VLP) packaging; upon transfection, Cas9 was cleaved by the protease from the viral pol domain in the host cell, leading to excellent gene editing efficiency (Mangeot et al., 2019; Hamilton et al., 2021; An et al., 2024; Hamilton et al., 2024). To enhance the delivery efficacy of VLP, the ratio between gag-POI and gag-pol should be optimized. Additionally, the cleavage linker between gag and POI is also crucial for transfection, as it should be exposed on the protein surface for effective recognition by protease (Banskota et al., 2022; Raguram A 2022; An et al., 2024). In addition to nuclease delivery by lentiviral-designed VLP, gag domain from human immunodeficiency virus (HIV), adeno-associated virus, Hepatitis B virus (HBV), Hepatitis C virus (HCV), or bacteriophages MS2, can also be assembled into VLP to carry other specific antigens to form VLP-based vaccines (Roldão A 2010; Klamp et al., 2011; Li et al., 2014). With sizes ranging from 20 to 200 nm, resembling pathogen-associated structural patterns, these VLP-based vaccines can efficiently elicit in vivo immune responses and have been approved for use in clinical trials(Roldão A 2010). Immune cells such as dendritic cells (DCs) and macrophages take up these vaccines through phagocytosis and micropinocytosis, and the present antigens from VLP to stimulate T cells for treating infectious diseases or cancer. However, viral-derived gag-pol in VLP, as a heterologous protein with strong immunogenicity, can also stimulate B cells to secrete neutralizing antibodies. These antibodies may remove VLPs from the body, thereby inhibiting the efficacy of VLP-based vaccines (Nooraei et al., 2021).

*PEG10* is a domesticated transposable element, which is expressed in mammals encoding two open read frames, PEG10-RF1(a gag-like protein) and PEG10-RF1/2 (a fusion of the gag and pol domains) (Lux et al., 2005). In brain, PEG10 has been identified as a component of stress granules and extracellular vesicles responsible for carrying mRNAs and proteins to transport information between cells (Pandya et al., 2021). Overexpression of PEG10 in HEK293FT cells resulted in the observation of extracellular vesicles containing the core capsid (CA) domain in the culture medium, demonstrating that endogenous retrovirus-like gag protein could be engineered into VLP for molecule transport (Segel M et al., 2021). Beside PEG10, the retrovirus-like protein Arc can also form VLP for cell communication, which is smaller than PEG10 (Ashley et al., 2018; Pastuzyn et al., 2018). Therefore, theoretically, endogenous retrovirus-like gag protein could be fused with antigens to produce VLP-based vaccines. However, endogenous VLP (eVLP) lack immunogenicity compared to viral VLP. This means that eVLP alone may struggle to activate dendritic cells (DCs) for inducing a robust immune response. To address this limitation, synthetic CpG oligonucleotide (CpG-ODN), a Toll-like receptor 9 (TLR9) agonist, has been explored as adjuvants in clinical vaccine trials due to their potent immunostimulatory properties(Coffman et al., 2010). DEC-205 is a multi-lectin receptor that has the binding capacity to CpG-ODN. For DCs, once CpG-ODN bonds to DEC-205, pinocytosis will be activated (Lahoud et al., 2012). Once CpG-ODN is connected to eVLP, the particles will further gain two advantages: first, CpG-ODN modified eVLP tend to bind DEC-205, for initiating DCs to engulf these particles; second, after endocytosis, the CpG-ODN can enhance DC maturation and antigen presentation as an adjuvant. Therefore, CpG-ODN modification represents an excellent strategy to improve immune responses for eVLP-based vaccines.

In order to overcome the challenges associated with peptide vaccine delivery and leverage the unique biological characteristics of PEG10, herein we firstly engineered PEG10 gag domain with liver cancer specific neoantigens to assemble into eVLP for peptide vaccine delivery. Furthermore, CpG-ODN was anchored on the surface of eVLP-antigen complex, named as ePAC to enhance the DC targeting efficacy and the antigen induced immune response. Afterwards, we further evaluated the antitumor effect of our ePAC in hepatocellular carcinoma (HCC) mouse model and confirmed that ePAC, as a cancer therapeutic vaccine, can activate immune response and regulate the tumor microenvironment. In general, our study reported a novel antigen delivery system which shows remarkable efficacy in transporting antigens to develop cancer therapeutic vaccine.

## Results

### Preparation and Characterization of PEG10-assembled eVLP

The protein transfection process relying on VLP produced by virus-associated gag domain is straightforward and highly efficient (Voelkel et al., 2010; Kaczmarczyk et al., 2011). As a type of retrovirus-like protein, mammalian-derived PEG10 possesses the canonical gag domain. We envision that the Protein of Interest (POI) could be fused to the PEG10 gag domain to create endogenous Virus-Like Particle (eVLP), which could then be delivered to targeted cells. To assess the transfection feasibility, eGFP was fused to the PEG10 gag domain behind the putative cutting site (NSQTD*PTEPV) (Lux et al., 2005) or a GS linker, along with the PEG10 gag-pol domain (Figure 1A). Plasmid VSVg is commonly used to assist retrovirus transfection. Therefore, as shown in Figure 1B, gag-eGFP or gag-pol-eGFP combining with VSVg were transfected to HEK293T cells for eVLP production. After 24 hours, the fluorescence from HEK293T cells were recorded (Figure. S1A). We observed that eGFP fused behind the gag domain showed superior expression level compared to the gag-pol fused structure. Therefore, we chose group 2 to further determine the secreted eVLP, which were collected from the supernatant of the transfected HEK293T cells and concentrated through ultracentrifugation. As show in Supplementary Figure 1B and 1C, gag-eGFP protein was detectable via western blot analysis in the concentrated supernatant, similar to its detection in cell lysate, with CD9 serving as the loading control (the maker of extracellular vesicles (Boker et al., 2018)). These results indicated that gag-eGFP could be self-assembled into eVLP for collection.

**Figure 1.**
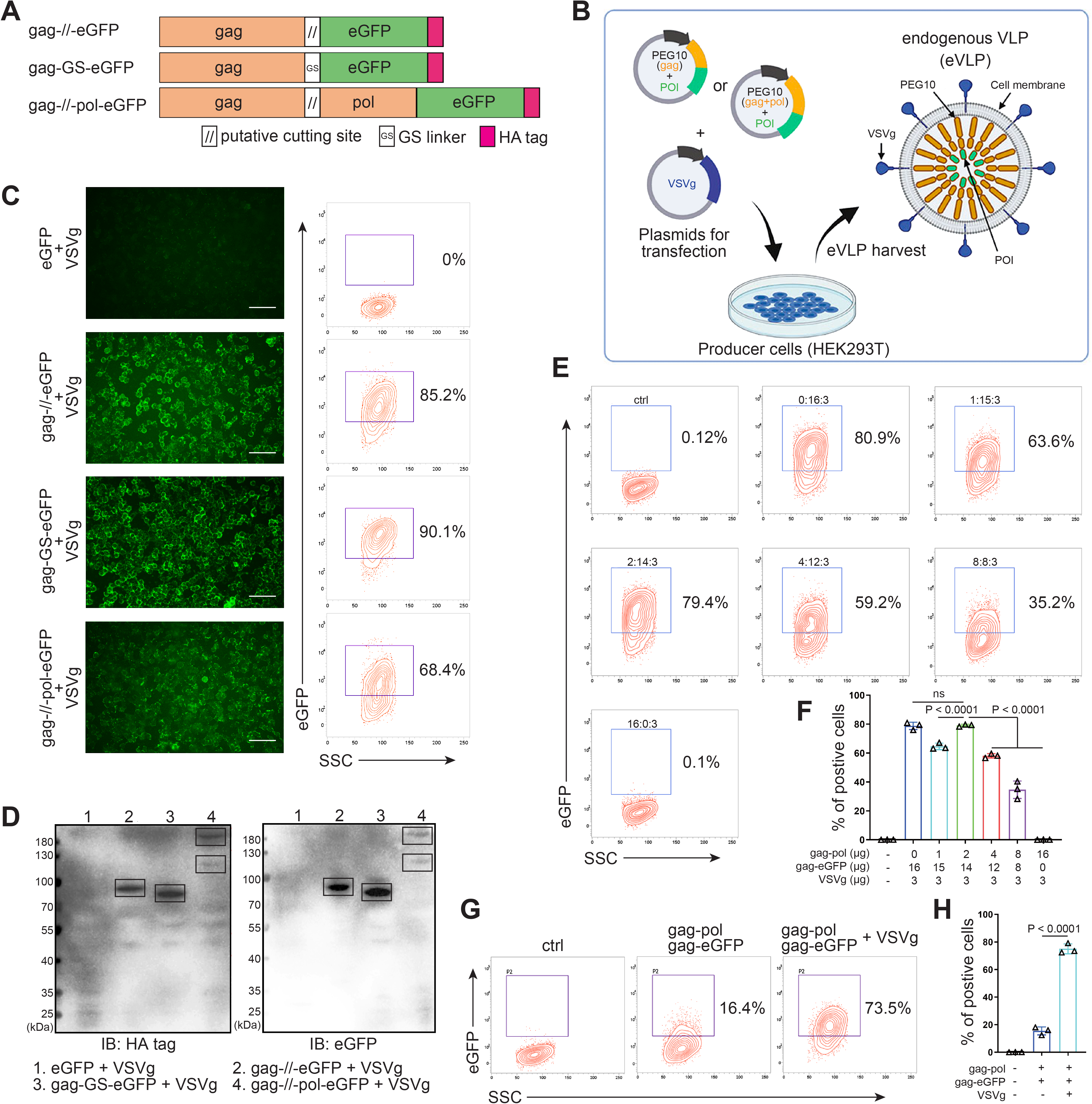
Designing and identifying the eVLP package system. (**A**) Schematic representation of PEG10 variants. The putative cutting site from PEG10 and GS linker were used to fuse PEG10 and eGFP together. HA tag was co-expressed in the C-terminal for subsequent identification. (**B**) The eVLP packaging process was illustrated in the schematic diagram. Plasmids from (A) were co-transfected with VSVg into HEK293T cells. eVLPs were harvested from the cell culture medium by ultracentrifugation. (**C**) The fluorescent image representing the eVLP transfected HEK293T cells (in blue) after 24h, and the positive percentage was analyzed by flow cytometry. Scale bar, 100Lμm. (**D**) The cells from (C) were harvested for HA tag and eGFP detection by western blot. 20 μl from each sample was loaded to the SDS-PAGE. Lane 1: eGFP, Lane 2: gag-//-eGFP, Lane 3: gag-GS-eGFP, Lane 4: gag-//-pol-eGFP (gag domain: ∼50 kD; gag-pol domain: ∼120 kD; eGFP: 27 kD). (**E**) and (**F**) Comparison of the eVLP transfection efficiency under different packaging strategies by flow cytometry. The plasmid ratio between gag-pol and gag-eGFP was optimized. The total plasmids for transfection in one 10-cm dish were 19 μg. (**G**) and (**H**) Identification of the auxiliary function of VSVg during eVLP transfection. Data are presented as the mean ± SEM.

Subsequently, the eVLPs collected from the four groups in Figure S1A were transfected into fresh HEK293T cells. Encouragingly, all gag and eGFP fused structure exhibited fluorescence in the cells, even with efficiency 90.1% by GS linker (Figure 1C). For cutting site connecting structure, it reduced to 85.2% possibly owing to the steric effect from the fused protein during gag domain accumulation in cellular membrane (Figure 1C). Meanwhile, the transfected HEK293T cells were collected and analyzed by Western Blot. The results further confirmed that eGFP can be transfected into HEK293T cells through PEG10 gag domain packaged eVLP (Figure 1D). Furthermore, when eGFP was fused to full-length PEG10, the fluorescence intensity was lower compared to the gag-only fusion type. But two distinct bands can still be observed during blotting, indicating successful self-cleavage by PEG10 protease domain (Figure 1C and 1D). These results pointed out that GS linker could assist gag domain accumulation after fusing POI, and co-packaging PEG10 pol domain could facilitate POI releasing from gag domain in the presence of cutting site. Thus, behind cutting sequence of PEG10 gag domain, we used GS linker to connect POI for following experiments, presenting as gag-POI.

Normally, gag-pol and gag-POI should be co-transfected to produce VLP. Increasing the percentage of gag-pol could make VLP more stable and release POI efficiently from gag domain, however, it also would reduce the loading capacity of VLP (Hamilton et al., 2021; Banskota et al., 2022). Thus, to carry sufficient targeting protein for transfection, the plasmid ratio between gag-pol and gag-POI should be coordinated. We then compared eVLP transfection efficiency to HEK293T cells with various ratios of gag-pol and gag-eGFP plasmids, and noticed that gag-pol 2 μg: gag-eGFP 14 μg: VSVg 3 μg was the most appropriate for targeting protein transfection (Figure 1E and 1F). Meanwhile, the function of VSVg for PEG10 based eVLP transfection was also determined. Without VSVg, the positive eGFP cells decreased dramatically from 73.5% to 16.4%, indicating that eVLP, like lentiviral or retroviral vectors, needed VSVg for transfection (Figure 1G and 1H). To sum up, we designed a new protein transfection strategy based on PEG10-packaged eVLP with high transfection efficiency.

### Envelope decoration of neoantigen-loaded eVLP

To prepare eVLP-based neoantigen vaccine, PEG10 gag domain was then fused with 7 tumor neoantigens from Hepa1-6 cell line (Chen et al., 2022) instead of eGFP, and the productivity of eVLP was evaluated by quantifying the total amount of proteins produced from one 10-cm dish of transfected HEK293T cells (eVLP vs control: 212±7.388 μg vs 76.59±7.463 μg, Figure S2). But in vivo, solely relying on the eVLP-antigen complex for antigen delivery to antigen-presenting cells (APCs) and immune response activation is not efficient. Although pseudotyped vectors expressing VSVg demonstrate robust mechanical stability and high titers for in vitro transduction, achieving effective transduction efficiencies for therapeutically relevant cells in vivo, such as hematopoietic stem cells, dendritic cells (DCs), and T cells, requires further modification of the vector surface by antibodies, peptides, or nucleotides (Lahoud et al., 2012; Lynn et al., 2020; Wei Y 2021). Here we utilized CpG-ODN to modify the eVLP-antigen complex, resulting in what we termed ePAC, which can interact with the DC surface protein DEC-205, thereby stimulating the intracellular receptor TLR9 and enhancing DC maturation. The modification process involved two steps: 1) The free amino group from the concentrated eVLP was reacted with NHS Ester carried by DBCO under 4L for 12h in PBS buffer. This step allowed for the anchoring of DBCO onto the eVLP. 2) CpG-ODN synthesized with 3’Azide (and 5’FAM) was added to DBCO anchored eVLP. This facilitated copper-free click chemistry, enabling the coupling of CpG-ODN to the eVLP (Figure 2A). Furthermore, by varying the concentration of DBCO-C6-NHS Ester from 0 to 14 nmol, ePAC exhibited different CpG-ODN loading efficiency as evidenced by agarose gel electrophoresis (Figure 2B and Figure S3). And the results showed that in a 200 µl eVLP reaction system, 3.5 nmol DBCO achieved the highest modification efficiency. Meanwhile, ePACs with elevated levels of CpG-ODN modification demonstrated robust fluorescent intensity in flow cytometry analysis (Figure 2C).

**Figure 2.**
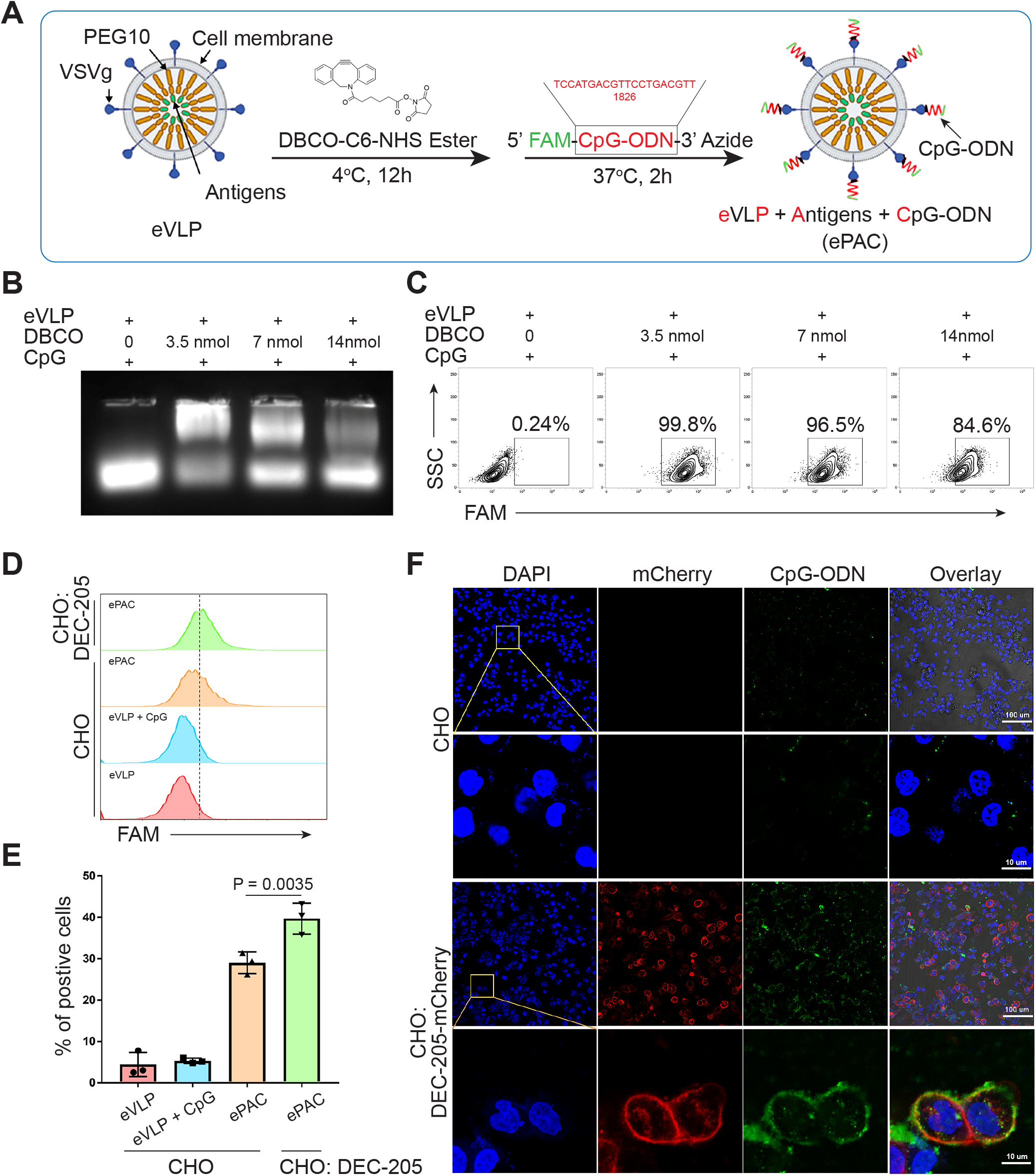
Envelope decoration of neoantigen-loaded eVLP. (**A**) Schematic of the decoration strategy for eVLP. DBCO-C6-NHS Ester was first anchored to eVLP under 4°C for 12h. Then, 5’-FAM-CpG-ODN-3’-Azide was added to trigger click reaction for 2h at 37°C. The CpG-ODN modified eVLP carrying antigens was named as ePAC. (**B**) Agarose gel electrophoresis of CpG-ODN modified eVLP with various DBCO-C6-NHS Ester concentration. eVLP: 200 μl per reaction, 5’-FAM-CpG-ODN-3’-Azide: 1 nmol per reaction. 20 μl sample was loaded in each lane. (**C**) The percentage of FAM positive HEK293T cells analyzed by flow cytometry. 50 μl modified eVLP from (B) was used to transfect HEK293T cells in a 24-well plate. (**D**) and (**E**) The bonding affinity between DEC-205 expressed CHO cells and ePAC was analyzed by flow cytometry (n=3 independent replicates; one-way ANOVA). CHO cells were transfected by the plasmid pCDH-DEC-205 to transiently express DEC-205. (**F**) The confocal microscopy images showing the bonding of DEC-205 and ePAC. CHO cells were transfected by lentivirus to express DEC-205-mCherry. Scale bar, 100Lμm. Data are presented as the mean±SEM. *p<0.05.

DEC-205 is a CpG-ODN receptor expressed by a variety of antigen presenting cells (Lahoud et al., 2012; Li et al., 2021). To explore the potential binding of CpG-ODN by DEC-205 and its impact on enhancing ePAC transfection efficiency, CHO cells, known for their ease of engineering to express large functional membrane proteins, were utilized to express mouse DEC-205. By flow cytometry analysis, ePAC transfection efficiency was 39.70±2.157 % in CHO cells with transient transfected DEC-205, which was significantly higher than that in wild type CHO cells (29.0 ± 1.514%, *p=0.0035*, Figure 2D and 2E). In addition to transient expression, DEC-205 fused with mCherry at the C-terminal was introduced into CHO cells via lentiviral transfection. The expression of DEC-205-mCherry was notably observable using mCherry fluorescence under confocal microscopy (Figure 2F). Meanwhile, following a 2-hour co-culture with ePAC, CHO cells expressing DEC-205-mCherry exhibited stronger FAM signal than that in wild type CHO cells (Figure 2F). These results suggested that CpG-ODN decoration could specifically improve the eVLP transfection efficiency in cells expressing DEC-205.

### In vitro stimulation of DCs by ePAC

The transfection efficiency of ePAC was further verified in DC2.4 cells and bone marrow-derived dendritic cells (DCs) from mouse (Figure 3A and 3B). With CpG-ODN modification, the transfection efficiency of eVLP for DC2.4 cells and DCs were 99.9% and 43.4%, respectively (Figure 3C and 3D). To confirm whether ePAC activated the intercellular receptor TLR9 signaling with cytokines secretion, the medium from DC2.4 cells were analyzed by ELISA. Compared to CpG-ODN or eVLP treated group, ePAC could significantly upregulate IL-6 and IL-12p70 secretion (Figure 3E). Furthermore, the DC2.4 cells were also collected for Western Blot to analyze TLR9-Myd88-P65 signaling. After 6h stimulation, ePAC treated group exhibited higher fold change ratio of phosphorylated P65 (0.79 vs 0.34) and Myd88 (0.63 vs 0.37) than eVLP group (Figure 3F). These results suggested that ePAC can be uptake by DC2.4 cells efficiently to activate the intracellular TLR9 signaling. To further investigate the immune response triggered by ePAC stimulated APCs in vitro, moused bone marrow derived DCs were co-culture with ePAC for two days. Subsequently, the expression levels of CD80 and CD86 (the hallmarks of DCs maturity) in CD11c^+^ DCs were measured by flow cytometry. In contrast to eVLP treated group, the ePAC treated group showed a significant increase in CD80 and CD86 expression (24.08±0.2355% vs 14.25±0.7854%, Figure 3G and 3H). Following this, mouse spleen-derived T cells were stimulated in vitro by the ePAC-transfected DCs for two days. On the third day, the fresh transfected DCs were reintroduced to the stimulated T cells for an additional two rounds of co-culture. Hepa1-6 cells were then co-cultured with the final activated T cells for 3 days to assess the killing efficiency mediated by DC-activated T cells. Post-co-culture, the cell culture supernatant was collected for cytotoxicity assays using LDH detection (Figure 3B).

**Figure 3.**
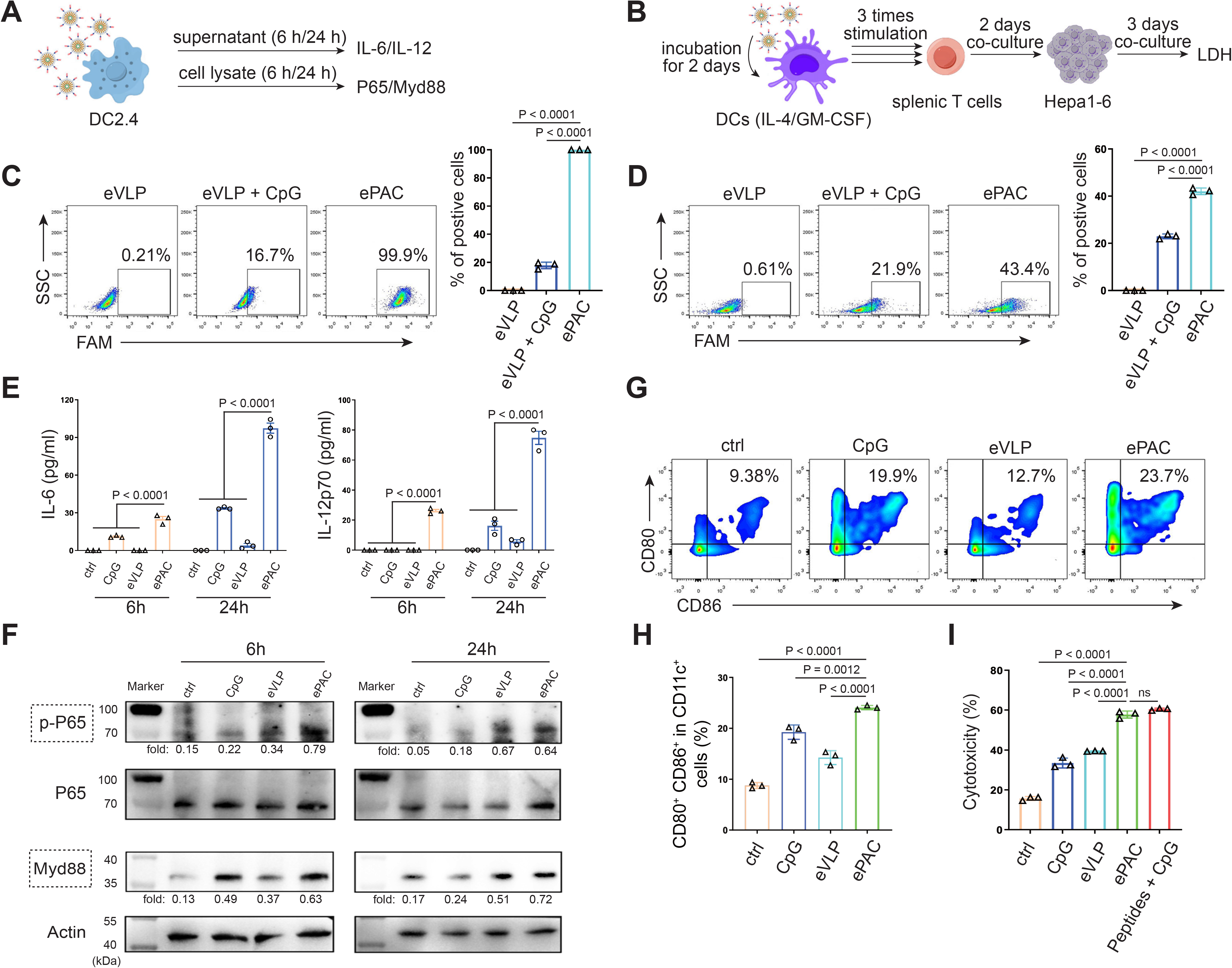
In vitro stimulation of DCs by ePAC. (**A**) The strategy for DC2.4 cell stimulation and detection. (**B**) The strategy for mouse-derived BMDC/T cell stimulation and Hepa1-6 co-culture. (**C**) The percentage and statistical analysis of ePAC transfection efficiency for DC2.4 cells by flow cytometry. (**D**) The percentage and statistical analysis of ePAC transfection efficiency for BMDCs by flow cytometry. **(E)** Cytokine secretion by DC2.4 cells at different time point after ePAC stimulation. **(F)** Western blot analysis of phospho-P65 and Myd88 expression at different time point after ePAC stimulation. P65 and Actin were used as loading control to calculate the fold change of p-P65 and Myd88 in different samples, respectively. P65: ∼65 kDa, Myd88: ∼33 kDa, Actin: 43 kDa. (**G**) and (**H**) The percentage and statistical analysis of matured DCs with CD80 and CD86 co-expression detected by flow cytometry (n=3 independent replicates; one-way ANOVA). (**I**) In vitro cytotoxicity against neoantigen-expressing Hepa1-6 cells determined by LDH assay (n=3 independent replicates; one-way ANOVA). Data are presented as the mean±SEM. *p<0.05, **p<0.01, ***p<0.001, ****p<0.0001; ns, no significance.

Notably, we found the percentage of Hepa1-6 cell death was dramatically increased in ePAC treated group (ePAC 57.73±1.018% vs eVLP 39.49±0.08%), similar with the positive control treated with Hepa1-6-derived neoantigen peptides plus CpG-ODN (Figure 3I). Together, our data suggested that ePAC is capable of delivering neoantigens to DCs for enhancing T cell-mediated antitumor activity.

### ePAC delivery and immune activation in vivo

The migration of ePAC to lymph nodes and its capacity to activate DCs are crucial determinants of the quality of the induced immune response. In order to investigate the accumulation of ePAC in lymph nodes, it was initially labeled with Dil and then subcutaneously injected into C57BL/6 mice (Figure 4A), while Dil and Dil plus eVLP (only carrying neoantigens) served as the control groups. As shown in Figure 4B, both ePAC and eVLP with the same vesicle structure produced from 293T cells, can efficiently accumulate in the draining lymph nodes two days post injection. The accumulation level of ePAC in lymph node is approximately 1.5 times higher than that of Dil plus eVLP group, and 5 times higher than that of Dil only group (Figure 4C). Subsequently, the collected lymph nodes were digested to prepare single-cell suspensions, and flow cytometry was employed to quantify the cellular uptake of ePAC in CD11c^+^ cells. As shown in Figure 4D and 4E, CpG-ODN modification enhanced the uptake efficiency of eVLP (ePAC 12.57±0.4055% vs eVLP 9.29±0.7225%, *p=0.0391*) compared to eVLP without CpG-ODN in vivo. To further determine the immune responses, the mice were subcutaneously injected with CpG-ODN, eVLP or ePAC for three times at different time points (Figure 4C). After sacrifice, the upregulation of MHC-II, CD80 and CD86 in CD11c^+^ DCs from lymph nodes was measured by flow cytometry. In comparison to eVLP treated group, the ePAC treated group exhibited significantly higher expression of MHC-II (ePAC 62.03±0.7753 vs eVLP 48.30±3.251, *p=0.0093*) and co-expression of CD80 and CD86 (ePAC 24.77±0.2333 vs eVLP 17.07±0.5239, *p=0.0023*) in CD11c^+^ DCs, indicating superior activation of DC maturation (Figure 4G-4J). Moreover, to analyze the formation of memory T cells induced by ePAC, the mouse splenic T cells were reactivated by DCs in vitro for interferon (IFN)-γ ELISPOT assay. The secretion of IFN-γ by T cells from ePAC stimulated group was approximately 2 times higher than that of eVLP treated group, (ePAC 156±13.45 vs eVLP 74±13.87, *p=0.0011*, Figure 4K and 4L). These results indicated that ePAC can be uptake by lymph nodes to activate DCs and T cells, thus holding significant potential as therapeutic vaccines.

**Figure 4.**
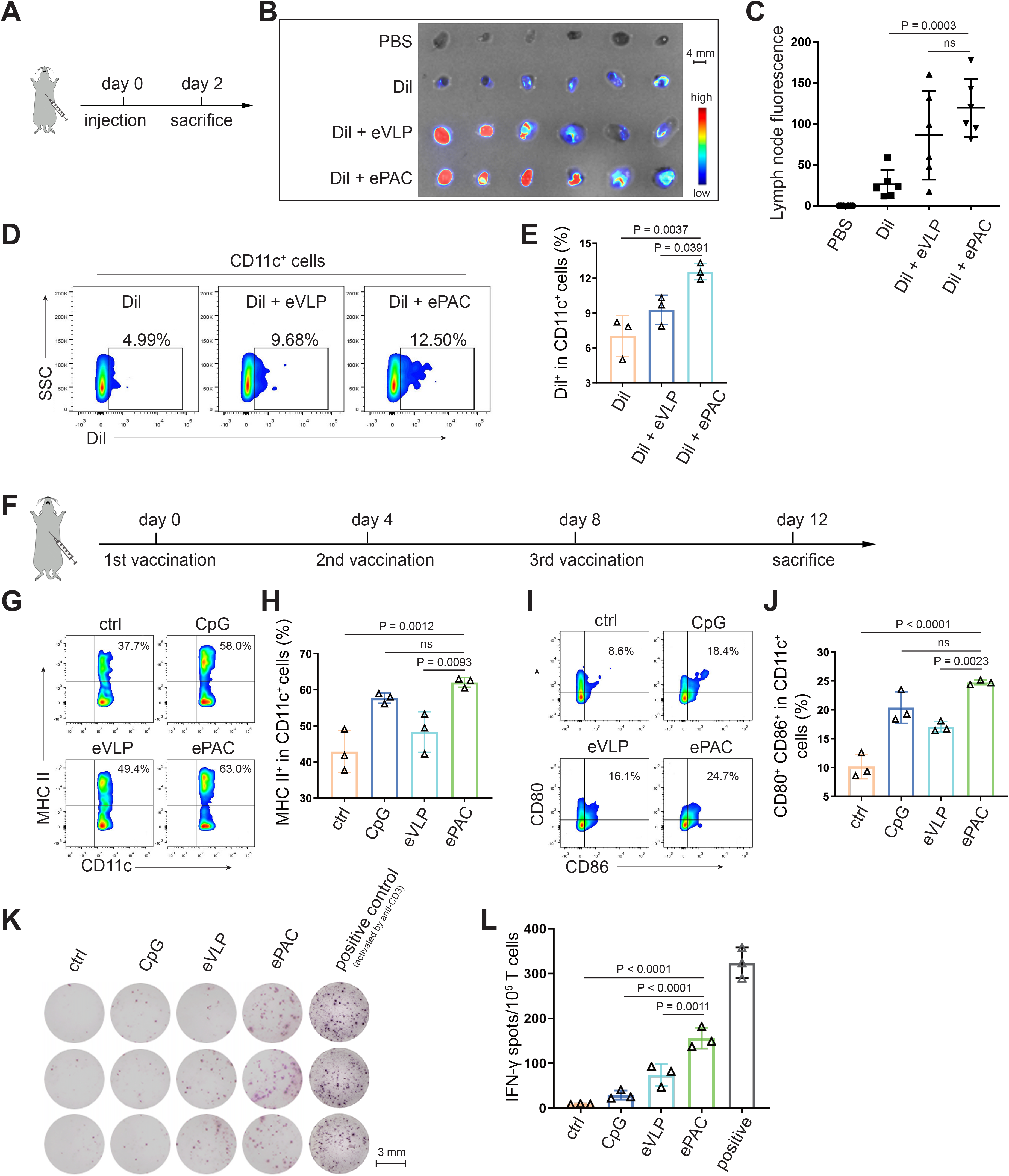
ePAC delivery and immune activation in vivo. (**A**) and (**B**) Ex vivo fluorescence image of isolated axillary lymph nodes (n=3 mice). eVLP was incubated with Dil for 6 h and then injected subcutaneously near the axilla of mice. 24 hours later, the axillary lymph nodes were collected and photographed. Every two lymph nodes came from one mouse. Scale bar, 4 mm (**C**) The statistical analysis of lymph node fluorescence intensity from (B) (n=3 mice; one-way ANOVA). (**D**) and (**E**) The percentage and statistical analysis of Dil positive DC cells in the lymph nodes detected by flow cytometry (n=3 mice; one-way ANOVA). (**F**) Schematic diagram of the vaccination protocol for C57BL/6 mice. (**G**) and (**H**) The percentage and statistical analysis of MHC expression level in the lymph nodes detected by flow cytometry (n=3 mice; one-way ANOVA). (**I**) and (**J**) The percentage and statistical analysis of matured DCs with CD80 and CD86 co-expression in the lymph nodes detected by flow cytometry (n=3 mice; one-way ANOVA). (**K**) and (**L**) ELISPOT assay showing neoantigen-specific reactivity of splenic T cells against Hepa1-6-derived neoantigens on day 12 after vaccination (n=3 mice; one-way ANOVA). The mouse anti-CD3 antibody was used to activate splenic T cells *in vitro* as the positive control for ELISPOT assay. Scale bar, 3 mm. Data are presented as the mean±SEM. *p<0.05, **p<0.01, ***p<0.001, ****p<0.0001; ns, no significance.

### Antitumor effect of ePAC in subcutaneous HCC model

The therapeutic efficacy of ePAC was assessed in C57BL/6 mice bearing Hepa1-6 tumor cells. A total of thirty mice with tumors of approximately 60 mm^3^ were randomly divided into five groups (n=6 per group). These groups were subcutaneously injected with PBS, CpG-ODN, eVLP, eVLP + CpG-ODN and ePAC, respectively, on the day 0, day 4, and day 8 (Figure 5A). Mice treated with PBS exhibited rapid tumor growth, whereas tumor growth in the other treatment groups significantly slowed down after three injections. Furthermore, ePAC with the ability to target DCs and increased stability by encapsulating peptides, exhibited significantly higher tumor growth inhibition efficiency (*p=0.0002*) comparing with the eVLP + CpG-ODN treated group similar to the simple mixture of neoantigen peptides and adjuvant (Figures 5B and 5C). Meanwhile, the Kaplan-Meier analysis of tumor progression free survival (PFS) also clearly demonstrated the therapeutic advantages of our ePAC (*p=0.0194*, Figure 5B). To further investigate whether ePAC exerted antitumor activity by activating immune responses, spleen and tumor samples were isolated from the treated mice. The percentage of effector memory T cells in the spleen was assessed via flow cytometry, revealing higher levels in the ePAC treated group (31.60±1.48%) compared to the eVLP group (14.22±1.377%) or the eVLP + CpG-ODN group (20.73±2.209%, Figure 5D and 5E). Within the tumors, the infiltration of CD4^+^ and CD8^+^ T cells were examined using immunohistochemistry. While changes in CD4^+^ T cells were not apparent across all groups, CD8^+^ T cell infiltration was notably stimulated by ePAC (Figure 5F and 5G). The activation of the infiltrating CD8^+^ T cells was further assessed by flow cytometry based on the expression level of 41-BB. As shown in Figure 5H and 5I, 4-1BB^+^ CD8^+^ T cells in total CD8^+^ T cells of ePAC treated group were 64.9%, substantially higher than in other groups (7.0% in PBS treated group, 18.4% in CpG-ODN treated group, 9.94% in eVLP treated group, 41.1% in eVLP + CpG-ODN treated group). Overall, these results suggest that the ePAC is an effective tumor vaccine capable of efficiently eliciting antitumor immunity.

**Figure 5.**
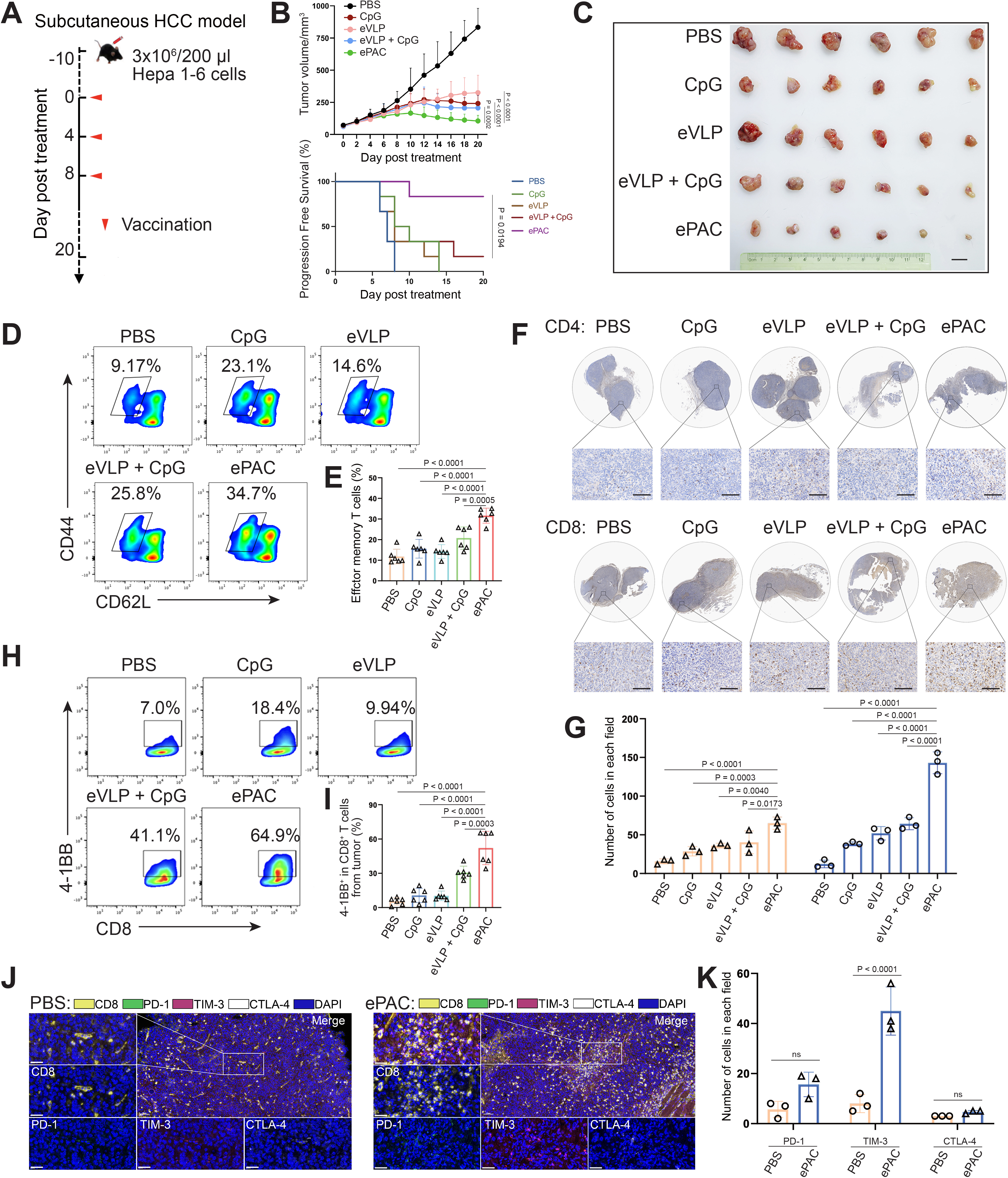
Antitumor effect of ePAC in subcutaneous HCC model. (**A**) Schematic diagram of vaccination-associated therapy in subcutaneous HCC model (n=6 mice). C57BL/6 mice were inoculated with Hepa1-6 cells on day −10 and immunized with the indicated vaccine formulations on day 0, 4 and 8 for a total of 3 treatments. (**B**) Growth curves of the tumor volumes in the indicated groups (n=6 mice; two-way ANOVA), and the progression free survival (PFS) of different treated groups was analyzed by Kaplan–Meier algorithm. The tumor reaching 200 mm^3^ was defined as progression. (**C**) Representative images of tumors harvested from tumor-bearing mice on day 20. Scale bar, 1 cm. (**D**) and (**E**) The percentage and statistical analysis of effector memory T cells in splenic CD8^+^ T cells detected by flow cytometry (n=6 mice; one-way ANOVA). (**F**) and (**G**) Immunohistochemical staining of CD4^+^ and CD8^+^ cells (brown) in the tumors collected at the dayL20 and the quantitation of CD4^+^ and CD8^+^ cells in each field (n=3, two-way ANOVA). Scale bar: 100Lμm in the lower panels. (**H**) and (**I**) The percentage and statistical analysis of activated T cells expressing 4-1BB from tumors detected by flow cytometry (n=6 mice; one-way ANOVA). (**J**) and (**K**) Representative images and the expression analysis of CD8, PD-1, TIM-3, and CTLA-4 in tumors by immunofluorescence (n=3; two-way ANOVA). Scale bar: 50Lμm. Data are presented as the mean±SEM. *p<0.05, **p<0.01, ***p<0.001, ****p<0.0001; ns, no significance.

Normally, strong activation of CD8^+^ T cells is often accompanied by high expression levels of immune checkpoints, which can inhibit their antitumor function. Next, we compared the expression level of PD-1, TIM-3 and CTLA-4 in CD8^+^ T cells from tumor tissues between PBS treated group and ePAC treated group. From the results of multicolor immunofluorescence, TIM-3 expression in ePAC treated group significantly increased (ePAC 45±5.568% vs PBS 8±2.082%, *p<0.0001*, Figure 5J and 5K). These results indicated that further combining ePAC with TIM-3 antibody (αTIM-3) might improve the antitumor efficiency.

### Enhancing ePAC antitumor efficacy in orthotopic HCC model by αTIM-3 combination

To accurately simulate the intrahepatic immune microenvironment and monitor vaccine response, we established an orthotopic hepatocellular carcinoma (HCC) model using Hepa1-6-luc-bearing mice. Given the upregulation of TIM-3 following ePAC vaccination, we aimed to enhance the antitumor immune response by combining ePAC with αTIM-3. The mice were randomly divided into 4 groups treated with PBS, αTIM-3, ePAC and ePAC + αTIM-3 (Figure 6A). As shown in Figure 6B, the combination of ePAC and αTIM-3 showed the lowest luciferase activity of tumors at the day 28, although the only αTIM-3 or ePAC treated mice showed reduced luciferase activity. Furthermore, the orthotopic tumor progress of 4/6 mice treated by ePAC + αTIM-3 was completely inhibited without any ghost, in contrast to the PBS (0/6), αTIM-3 (2/6), or ePAC treated alone (2/6) (Figure 6B). And the excellent synergistic therapeutic efficiency was further verified by the photograph of the liver and HE analysis (Figure 6C). To further investigate the changes in immune responses induced by the combination of ePAC and αTIM-3, we isolated the spleen and tumor from the treated mice. The effector memory CD8^+^ T cells in spleen and the infiltration of CD8^+^ T cells in tumor tissue were evaluated using flow cytometry. As shown in Figure 6D-6I, the percentages of effector memory CD8^+^ T cells (ePAC + αTIM-3 14.35±1.168% vs ePAC 11.32±0.6404% vs αTIM-3 11.28±0.3341%) and the infiltrated CD8^+^ T cells (ePAC + αTIM-3 55.72±2.733% vs ePAC 40.16±1.435% vs αTIM-3 37.79±2.278%) in mice treated with the combination of ePAC and αTIM-3 were higher than those in the other three groups . Additionally, the neoantigen-specific T cells from the infiltrated CD8^+^ T cells in tumor tissues collected from the combined therapy group were significantly higher than that in the other treated groups (ePAC + αTIM-3 9.06±1.258% vs ePAC 4.50±0.7195% vs αTIM-3 0.89±0.0746%). These results confirmed that our vaccine can activate the antigen-specific immune responses to efficiently elicit antitumor effects, highlighting the promising potential of combining ePAC with αTIM-3 as a therapeutic strategy for hepatocellular carcinoma.

**Figure 6.**
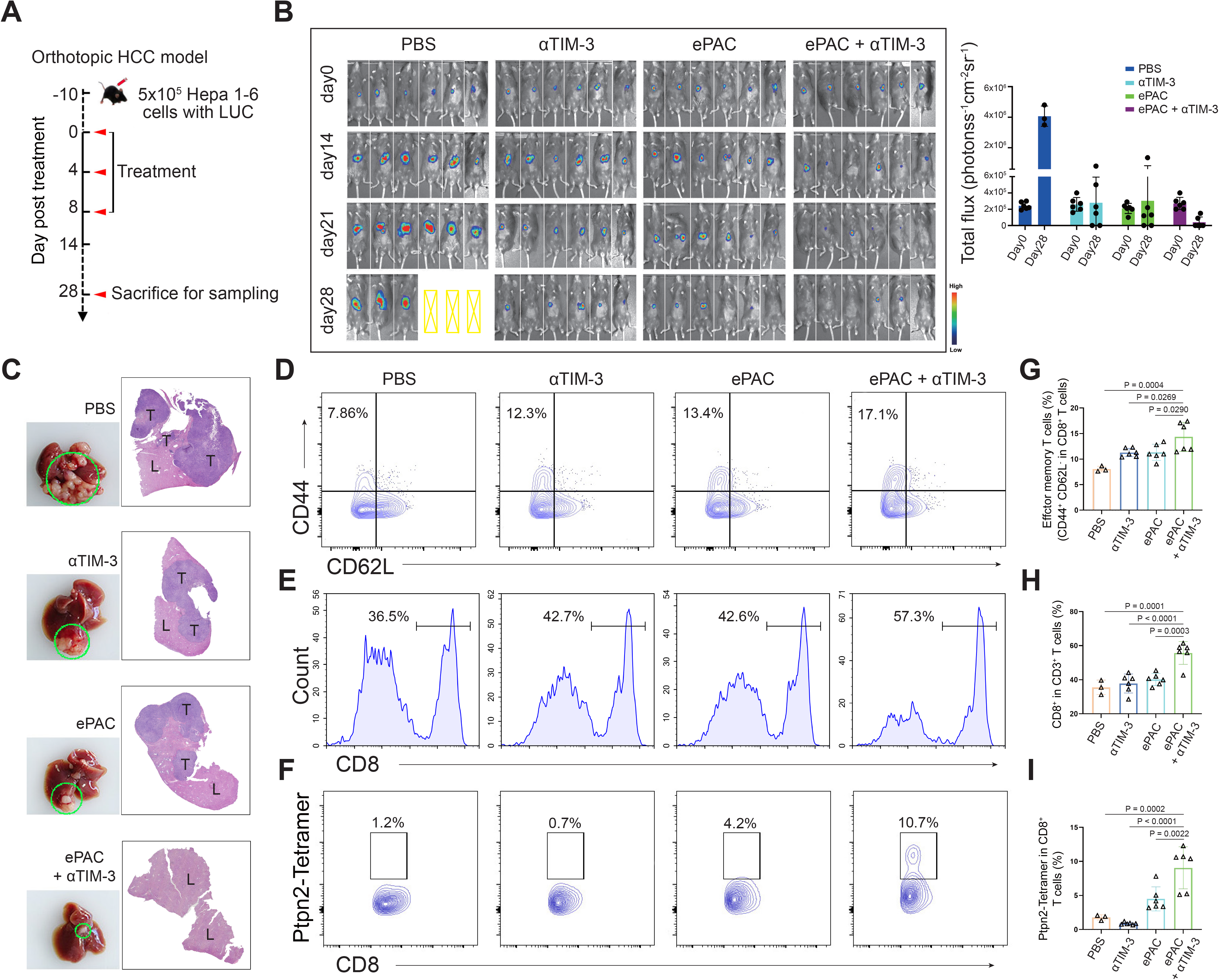
Evaluation ePAC antitumor efficacy in orthotopic HCC model by αTIM-3 combination. (**A**) Schematic diagram of vaccination-associated therapy in orthotopic HCC model (n=6 mice). The mice were sacrificed and sampled for analysis on the day of 28 after initiating treatment. (**B**) Tumor burden monitoring of PBS, αTIM-3 alone, ePAC alone, and ePAC plus αTIM-3 treated mice by bioluminescence imaging. (**C**) The photographs and H&E staining of tumor-bearing livers collected from mice after different treatments as indicated. “T” represents tumor tissues, “L” represents liver tissues, and there is the clear boundary with non-tumor sites. (**D-I**) Flow cytometry analysis of effector memory T cells in splenic CD8^+^ T cells (D and G), CD8^+^ T cell infiltration in the tumors (E and H), and Ptpn2_376-384_: H-2K^b^ specific CD8^+^ T cells in tumor infiltrating CD8^+^ T cells (F and I). Data are presented as the mean±SEM. *p<0.05, **p<0.01, ***p<0.001, ****p<0.0001.

### Antitumor effect by HLA-A*0201 restricted vaccine

To further confirm the function of T cell stimulation and antitumor effect of eVLP-packaged vaccine in human immune system, we incorporated an HBV-derived antigen (HBc18-27, with high binding affinity with HLA-A*0201) instead of previous neoantigens from Hepa 1-6 cells to PEG10 to prepare the ePAC vaccine. Peripheral blood mononuclear cells (PBMCs) from healthy donners with HLA-A*0201 restriction was used to derive monocytes, which were then induced with IL-4 and GM-CSF for 5 days to generate immature DCs (iDCs). Subsequently, the ePAC was added to stimulate iDC maturation for 2 days. After the stimulation, the DCs in ePAC treated group showed the highest level of maturation comparing to the eVLP treated group and control group (Figure S4), by using flow cytometry analysis. Then, the T cells from the same doner’s PBMCs with removing monocytes, were co-cultured with the stimulated DCs (Figure 7A). After three times of co-culture, the expression level of 4-1BB in T cells were significantly increased in ePAC treated group (ePAC 19.60±2.910% vs eVLP 8.367±2.485%, Figure 7B). To evaluate whether the DC-activated T cells led to killing of HBV-infected tumor cells, we performed tumor killing assays against HepG2.2.15 cells with stable expression of HBV antigens and HLA-A*0201 (Figure 7A). The activated T cells were then transfer to the plates seeded with HepG2.2.15 one day before at the ratio of 10: 1. The time dependent co-culture state was recorded by microscopy. From the image and LDH assay, we found that ePAC could ultimately improve T cell-mediated killing of tumor cells (ePAC 51.08±5.087% vs eVLP 27.56±2.433%, Figure 7C and 7D). These data suggested that human derived DCs could also be activated to present HLA-A*0201-restricted antigen by ePAC in vitro.

**Figure 7.**
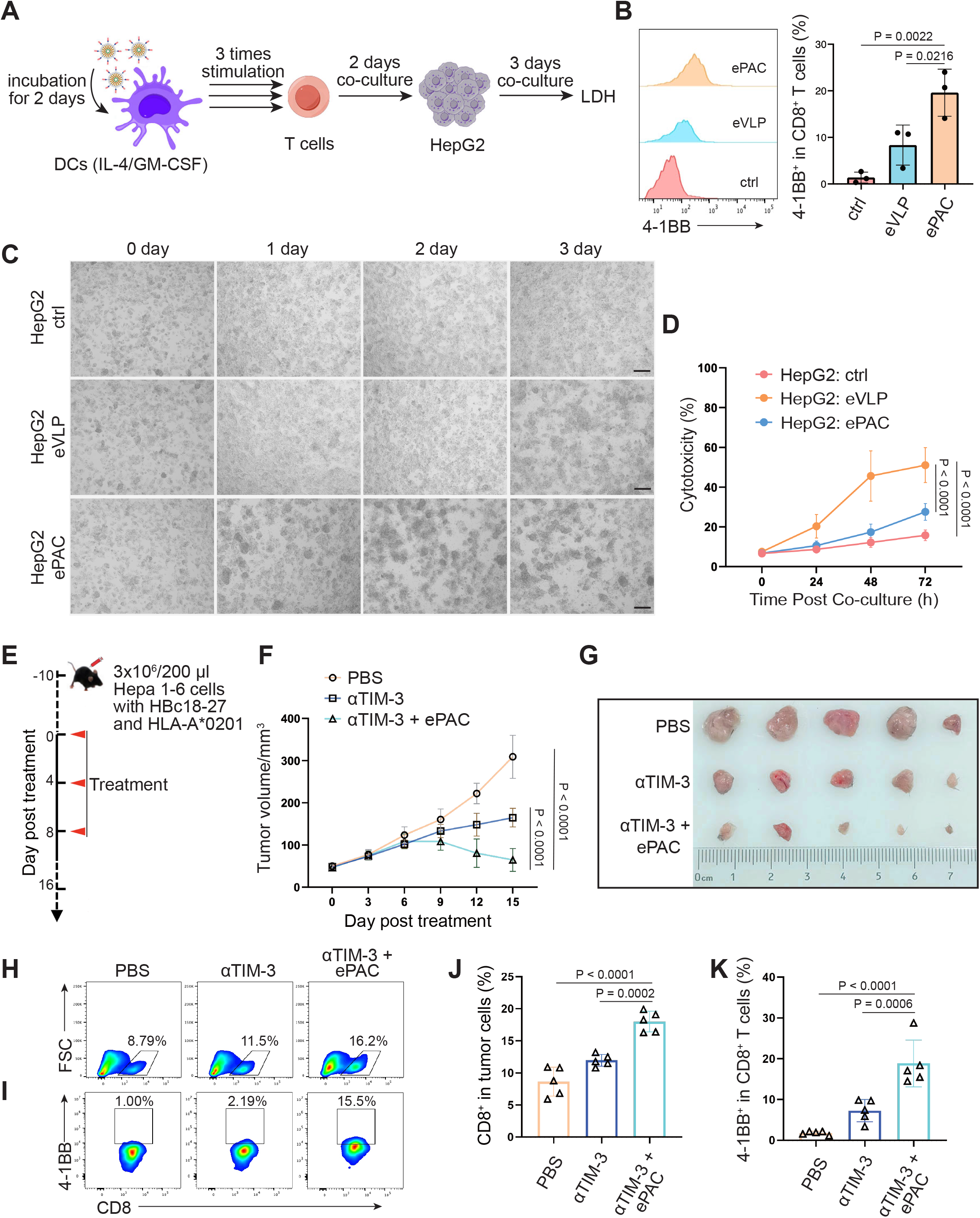
HBc18-27-specific immune response induced by eVLP-based vaccine. **(A)** The strategy for T cell stimulation and tumor cell killing. (**B**) The analysis of stimulated CD8^+^ T cells detected by flow cytometry (n=3; one-way ANOVA). PBMCs were first activated by anti-CD3 and anti-CD28 for 2 days and then co-cultured with ePAC stimulated DCs for three times. (**C**) Tumor killing assay of HepG2(2.15) cells visualized by microscopy. (**D**) In vitro cytotoxicity analysis after PBMC and HepG2 co-culture (n=3; two-way ANOVA). The supernatant from different time points were harvested for LDH assay. Scale bar, 100Lμm. (**E**) Schematic diagram of vaccination-associated therapy in subcutaneous HCC model constructed by HLA-A*0201 transgenic mice (n=5 mice). (**F**) Growth curves of the average tumor volumes in the indicated groups (n=5 mice; two-way ANOVA). (**G**) Representative images of tumors harvested from tumor-bearing mice on day 16. (**H-K**) Flow cytometry analysis of CD8^+^ T cell infiltration in the tumors (H and J) and the activated level of CD8^+^ T cells in tumors (I and K). n=6 mice; one-way ANOVA. Data are presented as the mean±SEM. *p<0.05, **p<0.01, ***p<0.001, ****p<0.0001.

The therapeutic efficacy of ePAC was further evaluated on HLA-A*0201 transgenic mice bearing Hepa1-6 tumor cells. Hepa1-6 cells were infected with adenovirus to express HLA-A0201 and the HBc18-27 epitope prior to seeding in mice (Figure S5A). 48 hours post-infection in vitro, strong mCherry signals were observed in the cells under an MOI of 500, and the HLA-A*0201 was consistently expressed on the cell surface (Figure S5B and S5C), suggesting that adenovirus-infected Hepa1-6 cells could potentially be targeted by ePAC vaccine-activated T cells in HLA-A0201 transgenic mice. After inoculating Hepa1-6 cells, a total of fifteen mice with tumors of approximately 60 mm^3^ were randomly divided into three groups and the time was recorded as day 0. These mice were subcutaneously injected with PBS, αTIM-3, and αTIM-3 + ePAC, respectively, on the day 0, day 4, and day 8 (Figure 7E). Two days before the treatment, the adenovirus carrying HBc18-27 and HLA-A*0201 were intratumorally injected to the tumors again to maintain the expression of HBc18-27 and HLA-A*0201. The mice inoculated with PBS and αTIM-3 suffered from continuous and rapid tumor growth, while in αTIM-3 + ePAC, tumor growth was significantly inhibited (Figure 7F and 7G). To further validate the immune response in tumors, the infiltrated CD8^+^ T cells in the tumors was analyzed by flow cytometry, revealing a higher percentage in total CD3 positive cells from the αTIM-3 + ePAC treated group (18.0±0.7197% vs 11.96±0.4130%) than that from the only αTIM-3 treated group (Figure 7H and 7J). Additionally, the activated level of the infiltrated CD8^+^ T cells were also tested based on the expression of 4-1BB. As shown in Figure 7I and 7K, there was a stronger activation level in the αTIM-3 + ePAC treated group (18.86±2.573%) compared to the αTIM-3 treated group (7.27±1.218%). Collectively, our data suggested that that ePAC, as tumor vaccine, can elicit antigen-specific immune responses in humanized mice with HLA-A*0201 restriction, highlighting its potential as an effective immunotherapeutic strategy for activating the human immune system to induce potent antitumor immunity.

## Discussion

Neoantigen vaccines designed by mixture of adjuvant and polypeptides have been demonstrated to effectively inhibit tumor growth and metastasis in mouse model (Mandelboim O 1995), and successfully treated several advanced solid tumors in clinical studies (Ott et al., 2017; Sahin et al., 2017). Our previous study also demonstrated that this type of personalized neoantigen vaccine can effectively prevent recurrence of primary liver cancer after surgery (Cai et al., 2021). All these studies have proved the antitumor effect by neoantigen-induced immune responses. Different from polypeptide vaccines, here we used PEG10 gag domain to directly fuse with neoantigens to package ePAC as cancer vaccine. This formulation has several advantages compared to polypeptides. Firstly, we can link multiple neoantigens into one plasmid for direct expression and harvest it as lentivirus by ultracentrifugation, so vaccines can be prepared easily in laboratory. Secondly, the cost of synthesizing polypeptides greatly saved, and it also avoids the situation that some polypeptides with strong immunogenicity, but high hydrophobicity cannot be synthesized. In addition, eVLP is more stable than polypeptide, which is essentially a vesicle produced from HEK293T cells. Although we did not compare the antitumor efficiency between polypeptide and eVLP in vivo, eVLP packaged cancer vaccine really induced neoantigen specific CD8^+^ T cells to antitumor. Furthermore, comparing to the virus-based delivery vectors, the lentiviruses although can stably integrate into the host genome but carry risks of insertional mutagenesis; adenoviruses although have high transduction efficiency but strong immunogenicity, which leads to fast clearance by the immune system of the host and affects the efficiency of the secondary injection. Instead, our VLPs offer low immunogenicity and superior safety, making them more suitable for repeated use and vaccine development. Therefore, we first demonstrate that PEG10 can co-express neoantigens and be packaged as eVLP-based cancer vaccine to induce strong immune response.

As a mammalian virus-like protein, PEG10 is the first to be developed to delivery mRNA and protein as eVLP (Abed et al., 2019; Pandya et al., 2021; Segel M et al., 2021). Arc is another endogenous virus-like protein with smaller amino acid length, mainly because it has lost functional protein domain during evolution and only retains the gag domain to support vesicle formation (Ashley et al., 2018; Pastuzyn et al., 2018). If Arc is engineered as eVLP, only two plasmids (gag-antigens and VSVg) are needed to prepare tumor vaccines, which could be simpler and more stable. But both PEG10 and Arc are mammalian homogenous proteins without the ability to induce host immune responses as natural viral proteins. And they also cannot specifically target to APC cells. In our study, CpG-ODN was anchored onto the surface of eVLP to activate DCs and direct eVLP to target DCs in vivo. In addition to serving as an adjuvant and targeting DCs, CpG-ODN possesses other advantages. Firstly, its nucleic acid sequence is short and easy to synthesize, resulting in low costs. Secondly, CpG-ODN as adjuvant has been approved by FDA. Furthermore, CpG-ODN is easy to modify with functional groups during synthesis, allowing for efficient conjugation to the surface of VLPs via click chemistry reactions.

Although neoantigen vaccines can induce immune responses to efficiently inhibit the development of tumors, some problems have been discovered in our previous studies. The most typical one is the high expressing level of PD-1 and TIM-3 in T cells after vaccination. When we used αPD-1 or αTIM-3 for combined intervention, the antitumor effect was superiors than neoantigen vaccine only (Chen et al., 2022; Zhao et al., 2022; Lin et al., 2024). In this study, we also measured the exhausted CD8^+^ T cells in subcutaneous tumors, and conducted the combined strategy with αTIM-3, which resulted in a more significant antitumor effect. Cytokines can also be used as a combined strategy for antitumor with neoantigen vaccines. IL-10 can reduce the exhaustion level of T cells through metabolic reprogramming (Guo et al., 2021; Zhao et al., 2024); IL-7 and IL-15 can moderately promote T cell proliferation and maintain the stemness of activated CD8^+^ T cells (Adachi et al., 2018; Silva et al., 2019). Therefore, in the next step, we will continue to explore some combined strategies based on ePAC vaccines and develop more efficient antitumor strategies for clinical translation.

In summary, we genetically engineered PEG10, one of the mammalian retrovirus-like protein, to establish an eVLP vector for antigen polypeptide delivery. Using this strategy, we further designed CpG-ODN modified tumor vaccine ePAC, which can effectively accumulate in the lymph nodes for DC targeting and stimulation. Thus, following the antigen presentation and cytokine secretion by the DCs, the activation of the tumor antigen-specific immune response leaded to considerable inhibition of tumor growth. These findings shed light on fundamental studies of fusion-based cargo delivery by eVLP, offering a promising avenue for the development of therapeutic vaccines targeting infectious diseases and cancers.

## Materials and methods

### Plasmids

PEG10 DNA sequence was synthesized according to pCMV-MmPeg10rc4 (addgene #174858). Then, the fragments of PEG10 including gag and gag-pol domain were amplified by PCR and inserted to pcDNA3.1 vector for plasmid construction. DEC205 was amplified by PCR from mouse bone marrow derived DCs, and mCherry was fused to the C-terminal of DEC205 and then inserted to pCDH vector for plasmid construction. Hepa1-6 cell derived seven neoantigens were identified in our previous study(Chen et al., 2022), and the DNA sequences were fused to the C-terminal of PEG10 gag domain cloned into pcDNA3.1 vector. The DNA sequence of HBc18-27 epitope was also synthesized and fused to PEG10. All DNA sequences were confirmed by sequencing after purification (EndoFree Maxi Plasmid Kit, TIANGEN Biotech). The sequences of these plasmids were listed in Table S1.

### Cell lines

HEK293T cells, CHO cells, DC2.4 cells and Murine HCC cell line Hepa1-6 cells were obtained from ATCC. HepG2.2.15 cells were purchased from IMMOCELL (Xiamen, China). All these cells were cultured in DMEM containing 10% FBS and 1 % penicillin/streptomycin at 37 °C in a humidified environment with 5% CO2. Luciferase-expressing Hepa1-6 (Hepa1-6-luc) cells were established through lentivirus transfection expressing luciferase reporter gene, and then the positive cells were selected by puromycin and maintained in low concentration of puromycin (Shanghai Genechem Co., Ltd).

### eVLPs production, purification and modification

eVLPs were produced by transient transfection of HEK293T cells. Briefly, HEK293T cells were seeded in 10-cm dish 24h before transfection. Then, the plasmids expressing PEG10 (16 μg) and VSVg (3 μg) were co-transfected per dish. For PEG10, it includes two plasmids, gag-POI and gag-pol. The ratio between gag-POI and gag-pol was regulated according to different experiments as indicated. After 48 h and 72 h, the cell culture medium was collected and centrifuged at 2000 g for 20 min to remove cell debris. The clarified eVLP-containing supernatant was then filtered by a 0.45-μm PVDF filter (EMD Millipore, #SE1M003M00), and ultracentrifuged at 120,000 g for 1.5h at 4°C in a Beckman Coulter Optima XPN-80 ultracentrifuge. Afterwards, the supernatant was discarded, and the pellet that finally harvested from one 10-cm dish around 20 ml cell culture medium was then further resuspended by 200 μl PBS. For eVLP that were used in the following in vitro experiments, it was concentrated 100-fold from cell culture medium using Lenti-X™ Concentrator (Takara, #631231) according to the manufacturer’s protocol. For eVLP that were injected into mice, it was concentrated 1000-fold by ultracentrifugation using a cushion of 20% (w/v) sucrose in PBS. To modify the concentrated eVLP by CpG-ODN, the DBCO-C6-NHS Ester (30 mg/ml, MACKLIN, China) was added into eVLP overnight at 4°C, then dialyzed using an ultra-centrifugal filter tube (Amicon® Ultra-0.5 50k device, Millipore, Darmstadt) extensively with PBS for three times. Afterwards, the CpG-ODN-Azide (100 μM, General Biol., China) was then further reacted with DBCO-eVLP through click chemistry at 37°C for 2h. Subsequently, the product was dialyzed again using an ultra-centrifugal filter tube against PBS for three times. The CpG-ODN modified eVLP was stocked at −80°C for further usage.

### Adenovirus production

Adenovirus vectors were produced by transfecting HEK293A cells with transgenic transfer plasmid pDC315 and pBHGlox(delta)E1, 3Cre packaging plasmid in a 1:1 molar ratio, normalized to a total mass of 20 μg plasmid/10-cm dish of cells, incubated with 60 μl PEI in 500 μl Opti-MEM medium. After two weeks, the transfected cells were harvested using 1 mL PBS and lysed through three cycles of repeated freeze-thaw (−80°C/37°C). Then, the cell lysate was centrifuged at 5000 g for 10 minutes, followed by filtration using a 0.45-μm PVDF filter (EMD Millipore, #SE1M003M00), resulting in the collection of the first generation of adenovirus. To amplify the adenovirus, the first generation was seeded into fresh HEK293A cells for 3 days. Subsequently, the cells were harvested and lysed as described above to collect the second generation of adenovirus, which can be further amplified and used to transfect target cell for gene expression.

### Western blot

Cell or eVLP lysates were prepared in RIPA lysis buffer (Beyotime Biotechnology, China) containing PMSF and a protease inhibitor cocktail (MedChemExpress, China). After centrifugation, protein samples were subjected to 10% SDS-PAGE and transferred onto polyvinylidene difluoride (PVDF) membranes (Roche, Mannheim, Germany). Afterwards, the membranes were blocked in TBST (1 mM Tris–HCl, pH 7.4, 150 mM NaCl, 0.05% Tween-20) containing 5% BSA for 1 h and subsequently incubated overnight at 4°C with diluted primary antibodies against HA tag, GFP, P65, Phospho-P65, Myd88 and Actin. Information on antibodies used was given in Table S2. The band was quantified using the FluorChem FC2 system (Alpha Innotech Corporation St. Leonardo, CA, USA).

### In vivo antitumor efficacy evaluation

Female C57BL/6 mice (6–8 weeks old) were purchased from China Wushi (Shanghai, China). HLA-A*0201/K^b^ transgenic mice were purchased from National Human Disease Animal Model Resource Center (Beijing, China), which are a well-established model for the study of HLA-A*0201 restricted epitopes and vaccine development (Lorena Passoni 2002; Gritzapis et al., 2004; Sun et al., 2014). All animal experiments were approved by the Animal Ethics Committee of Mengchao Hepatobiliary Hospital of Fujian Medical University. The subcutaneous HCC model was established by subcutaneous injection of 3×10^6^ Hepa1-6 cells into the right axilla. The orthotopic HCC mouse model was established as follows: 1) the mice were intraperitoneally anesthetized with 50mg/kg of pentobarbital; 2) the skin was prepared aseptically, and the midline laparotomy was performed after the abdomen shaving; 3) liver subcapsular inoculation with 5×10^5^ Hepa1-6-luc tumor cells mixed with the matrigel plugs; 4) the abdominal muscle and skin were fully-layer sutured.

Tumor monitoring of subcutaneous HCC cancer model was performed by a single operator every two days in two dimensions using calipers. Tumor volume was calculated from the measured data as follows: V=AB^2^/2 (A is the long diameter and B is the short diameter). When the tumor volume grew to 50-80 mm^3^, it was recorded as day 0, and then the mice were randomly divided into 5 groups (n=6) to start treatment. In the experimental group, CpG-ODN (10 μl/mouse, from 100 μM stock), eVLP (200 μg/mouse), and ePAC were administered subcutaneously to each tumor-bearing mouse on day 0 day 4 and day 8, respectively. The control group was treated for the same duration with PBS injection. For the orthotopic HCC cancer model, the mice were randomly divided into 4 groups (n=6): PBS, αTIM-3 (100 µg/mouse, Leinco Technologies, USA), ePAC (200 μg/mouse), ePAC (200 µg/mouse) + αTIM-3 (100 µg/mouse) after 10 days of tumor inoculation (recorded as day 0) and then the treatment was initiated. At the end of treatment, all mice were sacrificed by cervical dislocation and the number of tumor nodules was recorded. For the subcutaneous HCC model constructed by HLA-A*0201/K^b^ transgenic mice, the Hepa1-6 cells were infected by adenovirus to express HBc18-27 epitope and HLA-A*0201 before subcutaneous injection. Tumor monitoring was performed every three days and tumor volume was calculated from the measured data as previously outlined. The tumor volume grew to 50-80 mm^3^ was recorded as day 0, then the mice were randomly divided into three groups (n=5): PBS, αTIM-3 (100 µg/mouse) and αTIM-3 (100 µg/mouse) + ePAC (200 µg/mouse) to start treatment.

### Flow cytometry

Cells were harvested by 1.5 mL tubes and washed by 300 µl PBS. Then the samples were stained with the antibodies as needed for 30 min under dark. The detailed information on antibodies including anti-CD11c-APC, anti-CD80-PE, anti-CD86-PeCy7, anti-MHC-II-PE, anti-CD3-FITC, anti-CD8-APC, anti-CD44-PeCy7, and anti-CD62L-PerCP was given in Table S2. Ptpn2-Tetramer-PE was purchased from BioReagent Unit of Cancer Research Center of Xiamen University (Xiamen, China). After staining, the samples were washed two times by PBS and finally resuspended by 300 µl PBS for detection. Flow cytometry was carried out on the FACScalibur flow cytometer (BD, San Diego, USA). Data were analyzed using FlowJo software (Treestar, Inc., San Carlos, CA, USA).

### Cell imaging with confocal

The lentivirus carrying DEC205-mCherry was produced by four plasmids system (pMDL: 3.3 ug, pREV: 3.3 ug, VSVg: 3.3 ug, pCDH-CMV-DEC205-mCherry: 10 ug for one 10-cm dish transfection) and purified similarly to eVLP by ultracentrifugation from cell culture medium. CHO cells were transfected by this lentivirus to express DEC205-mCherry for three days and positively selected by puromycin. After two hours co-culture with ePAC, CHO cells were washed by PBS for three times and imaged via confocal microscopy (CLSM) using an LSM 780 instrument from Germany. The nucleus was stained with DAPI (excited at 405 nm), while the ePAC with FAM modification was excited by 488 nm and mCherry was excited at 561 nm.

### Enzyme-linked immunosorbent assay

The supernatant from DC2.4 cells stimulated by ePAC was collected and centrifuged at 2000 g for 10 min to remove the cell debris. Mouse IL-6 and IL-12p70 were quantified using Elisa kits (Boster Biological Technology, China) following the manufacturer’s protocols.

### Immunogenicity validation by Elispot

CpG-ODN, eVLP and ePAC were injected subcutaneously in the lateral flank of C57BL/6 mice on days 0, 4 and 8, respectively. The dose usage for each mouse was similar to the experiments of in vivo antitumor efficacy evaluation. The mice were sacrificed on day 12 and the splenic T cells were harvested for ex vivo interferon (IFN)-γ ELISPOT assay. IFN-γ secretion of mouse splenic T cells were detected by ELISPOT kit (Mabtech, 3321-4APT-10). Briefly, bone marrow derived-DCs (BMDCs) were isolated from 6-8 weeks old naive C57BL/6 mouse femurs and tibias. After removal of residual soft tissue and epiphyses, the marrow was collected by flushing the canals with PBS and centrifuged at 800g for 5 min at room temperature. The precipitated cells were lysed with 1mL Red Blood Cell Lysis Solution (Gibco) for 4 minutes, centrifuged at 800g for 5 minutes, and washed twice with PBS. At day 0, 2 million cells/well were added into a 6-well plate and cultured with 2 mL RPMI-1640 medium/well (10 ng/mL mIL-4, and 20 ng/mL mGM-CSF, GenScript, China) to obtain BMDCs at 37°C with 5% CO_2_. Half volume of medium was changed at day 3. At day 6, the BMDCs were pulsed with CpG-ODN, eVLP or ePAC for 48h. For ELISPOT assay, 2×10^4^ stimulated BMDCs were co-incubated with 2×10^5^ splenic T cells in a multiscreen 96-well filtration plate (Mabtech, 3321-4APT-2) at 37L with 5% CO_2_ for another 48h. BMDCs pulsed with PBS were used as the negative control. Then, the plates were washed and subsequently incubated with detection antibody (R4-6A2-biotin, 1 µg/mL, 100 µl/well) for 2 hours at room temperature. Afterwards, the plates were washed again and then incubated with Streptavidin ALP (1:1000 dilution, 100 µl per well) for 1 hour at room temperature. Subsequently, 3, 3’, 5, 5’-Tetramethylbenzidine (TMB) substrate solution was added to each well and incubated for 4-8 min at room temperature before adding deionized water to stop the reaction. Finally, IFN-γ spot-forming cells were imaged and analyzed by ELISPOT Analysis System (AT-Spot-2200, Beijing Antai Yongxin Medical Technology, China).

### In vitro Cytotoxicity Assays

The cell culture medium from different co-culture system was collected and centrifuged at 250 g for 10 min. The supernatant was further diluted for LDH detection according to the product manual (MK401, Takara). Briefly, 100 μl sample was transferred into a 96-well flat bottom plate following 100 μl reaction mixture. Then, the plate was incubated at room temperature for 30 min protected from light. Finally, the absorbance of the samples at 490nm was measured. The cytotoxicity was calculated by the equation: Cytotoxicity = (Absorbance (sample) - Absorbance (low control)) / (Absorbance (high control) - Absorbance (low control)) * 100%

### HE and Immunohistochemistry staining

HE staining was performed according to routine protocols. Briefly, after finishing deparaffinization and rehydration, the organ sections were stained 5 min with hematoxylin solution (G1004, Servicebio, China), followed by 5Ldips in 1% acid ethanol (1% HCl in 75% ethanol) and then rinsed in distilled water. Then the sections were stained with eosin solution (G1001, Servicebio, China) for 3Lminutes and followed by dehydration with graded alcohol and then clearing in xylene. The HE-stained slides were digitized, and the crop images were collected.

For immunohistochemistry staining, the paraffin-embedded tumor sections were mounted on slides, dewaxed in xylene, and rehydrated in graded alcohol washes.

Afterwards, the slides were heated by microwave in 0.01 mol/L trisodium citrate buffer for antigen retrieval. Then, 5% BSA was used to block nonspecific bonding sites for 30 minutes and 3% H_2_O_2_ was used to suppress endogenous peroxidase activity. The slides were then incubated with primary mouse antibody (CD4 and CD8, detailed information from Table S2) overnight at 4°C and washed with Tris-buffered saline (TBS) before incubation with labeled polymer horseradish peroxidase rabbit antibody for 30 minutes. Counterstaining was performed with hematoxylin. Finally, the slides were dehydrated through ascending alcohols to xylene and mounted to take photos. The index of CD4 and CD8 was given as the ratio between positive nuclei and total number of nuclei per visual field. Three visual fields per sample were randomly selected for analysis.

### Immunofluorescence staining

The primary antibodies for the immune inhibitory receptor panel included anti-CD8 (ab217344, Abcam), anti-PD-1 (ab214421, Abcam), anti-TIM-3 (ab241332, Abcam), and anti-CTLA-4 (ab237712, Abcam). Nuclei were highlighted using DAPI. The primary antibodies were added after successful blocking and incubated for one hour at room temperature, followed by secondary antibody incubation for 30 minutes at room temperature. The images with the final fluorescent were taken by the PhenoImager® HT instrument (Akoya Biosciences, USA)

## Data analysis

Statistical analysis of data was analyzed through one-way or two-way analysis of variance (ANOVA) for comparison among multiple groups. *p<0.05 was set as statistically significant. **p<0.01, ***p<0.001, ****p<0.0001. Results are presented as mean ± standard error of the mean (SEM) at least three experiments by GraphPad Prism software (San Diego, USA).

## Data availability

The data used to generate the results are shown in Figures and Supplementary Figures are available as supplementary information. Source data are provided with this paper.

## Ethic Statement

The studies involving human-derived blood were conducted in compliance with ethical regulations and approved by the Ethics Review Committee of Mengchao Hepatobiliary Hospital of Fujian Medical University (KESHEN 2021_100_02). All animal procedures were conducted according to the “National animal management regulations of China” and approved by the Animal Ethics Committee of Mengchao Hepatobiliary Hospital of Fujian Medical University.

## Acknowledgements

This work was supported by the National Natural Science Foundation of China (no. 82202027 and U22A20328); the Major Research Projects for Young and Middle-aged Talent of Fujian Provincial Health Commission (2022ZQNZD014), the Young and Middle-aged Talent Training Project of Fujian Provincial Health Commission (no. 2022GGA049); the Scientific Foundation of Fujian Province (no. 2023J06049 and 2022J011279); the Startup Fund for Scientific Research, Fujian Medical University (no. 2021QH1157).

## Author contributions

R.T., Y.Z., Z.C., and X.L. designed the study. R.T., L.G., T.W., T.C., H.Y., H.Y., and F.L. conducted the experiments. R.T., L.G., T.C., Z.C., and X.L. analyzed and discussed data. R.T., Z.C., and X.L. wrote and reviewed the manuscript. Z.C., and X.L. supervised the study. All authors read and approved the final manuscript.

## Declaration of interests

The authors declare no competing interests.

## Notes

### Competing Interest Statement

The authors have declared no competing interest.

### Summary of Updates

1. Section on Results of Envelope decoration of neoantigen-loaded eVLP updated to clarify the modification efficiency when the DBCO concentration is lower than 3.5 nmol. 2. Section on Results of Antitumor effect of ePAC in subcutaneous HCC model updated to clarify the benefit of using the compact vector other than just free peptide and CpG. 3. Section on Results of Antitumor effect by HLA-A*0201 restricted vaccine updated to clarify whether human DC cells can be activated by ePAC in vitro. 4. Section on Discussion updated to clarify the advantages of low immunogenicity viruses as vaccines compared with conventional adenovirus and lentivirus. 5. Figure 1, 2, 3, 4, 5 and 6 revised; 6. Supplemental files updated.

