## Supplemental Information for "Engineering PEG10 assembled endogenous virus-like particles with genetically encoded neoantigen peptides for cancer vaccination"

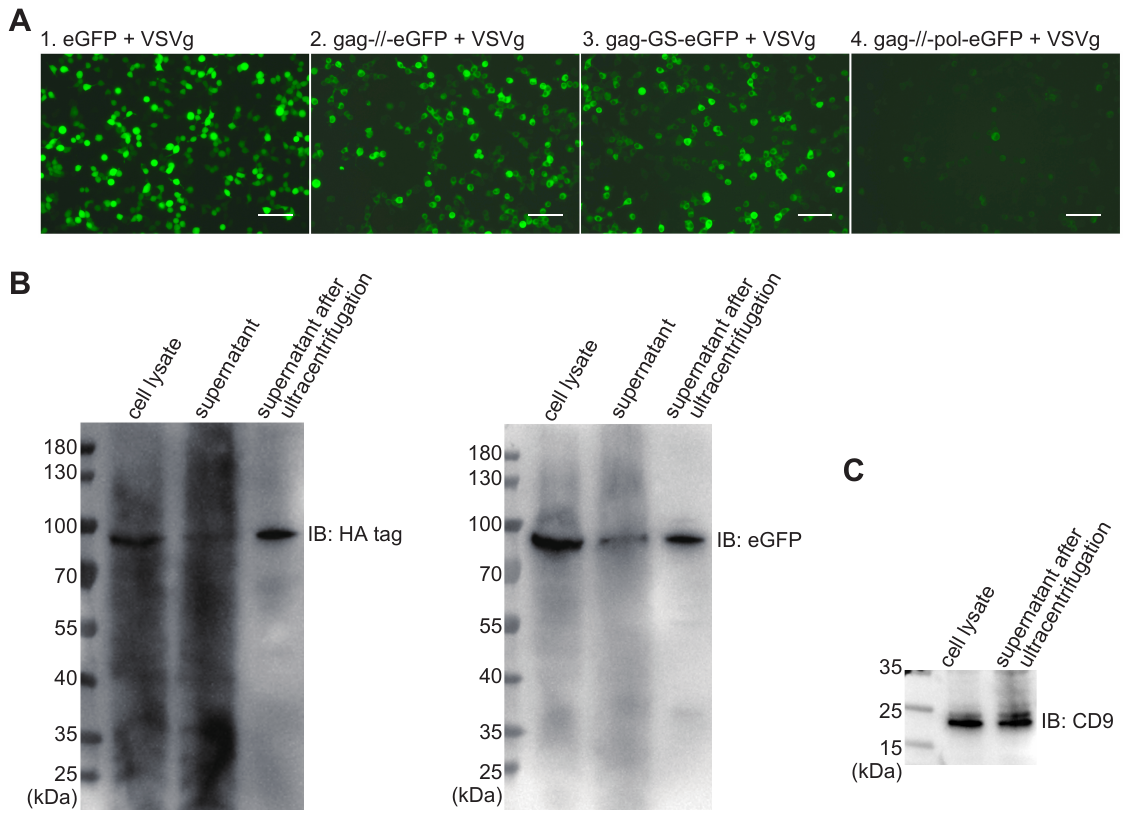

**Supplementary Figure 1. Evaluation of plasmid expression**

(A) The fluorescent images after 24 hours expression by plasmid transfection in HEK293T cells from the Fig. 1A. For transfection in one 10-cm dish, (gag plus) eGFP was 16 μg and VSVg was 3 μg. Scale bar, 100 μm. (B) Western blots for gag-//-eGFP expression after 3 days in HEK293T cells from (A). The concentration of eVLP from supernatant after ultracentrifugation was 100-fold higher than that before. (C) Western blot for CD9 (22 kDa).

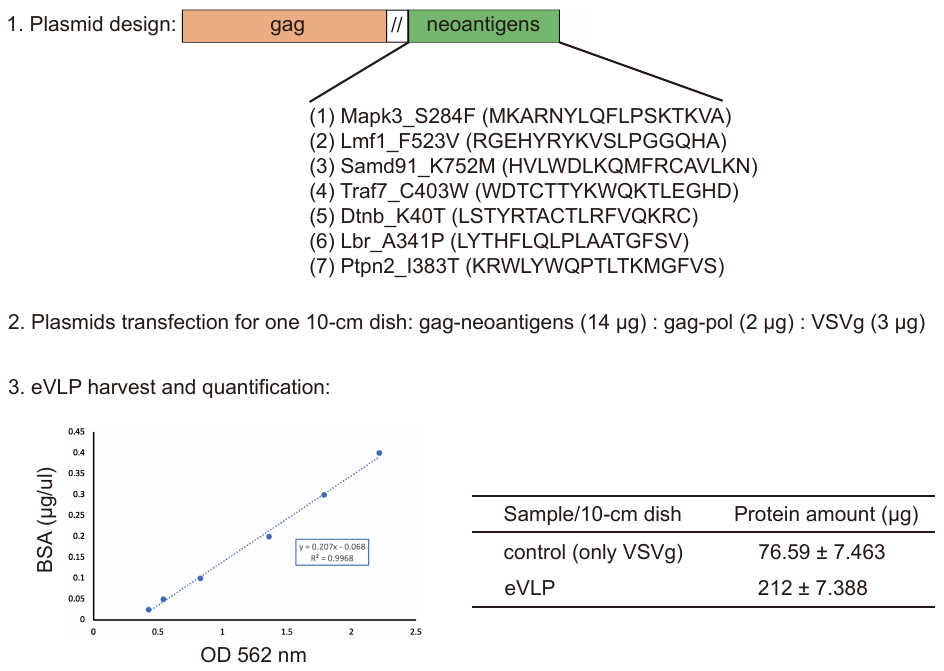

**Supplementary Figure 2. The eVLP-based neoantigen vaccine preparation**

The neoantigens (Mapk3_S284F, Lmf1_F523V, Samd91_K752M, Traf7_C403W, Dtnb_K40T, Lbr_A341P, Ptpn2_I383T from Hepa1-6 cell line) were fused together by GGGS, and then were connected to gag domain C-terminal for eVLP production. Three plasmids including gag-neoantigens, gag-pol and VSVg, were co-transected in HEK293T cells. The supernatant containing eVLP was harvested and concentrated by ultracentrifugation. The total proteins from the concentrated samples were finally determined by BCA.

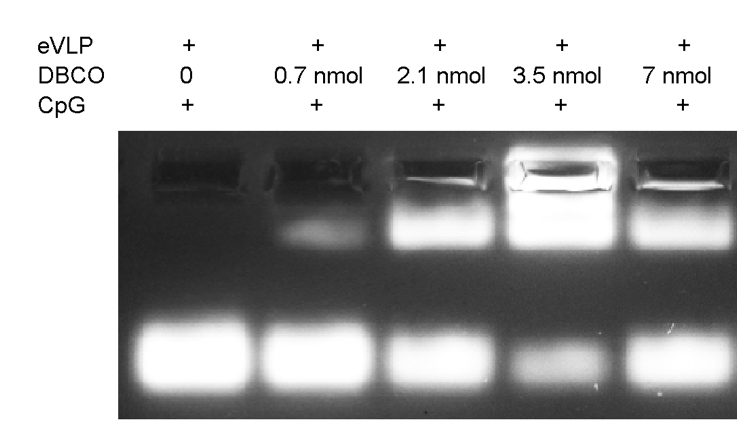

**Supplementary Figure 3. Agarose gel electrophoresis of CpG-ODN modified eVLP with various** **DBCO-C6-NHS Ester concentration.**

The concentration of DBCO-C6-NHS Ester included 0, 0.7, 2.1, 3.5 and 7 nmol. eVLP: 200 μl per reaction, 5’-FAM-CpG-ODN-3’-Azide: 1 nmol per reaction. 20 μl sample was loaded in each lane.

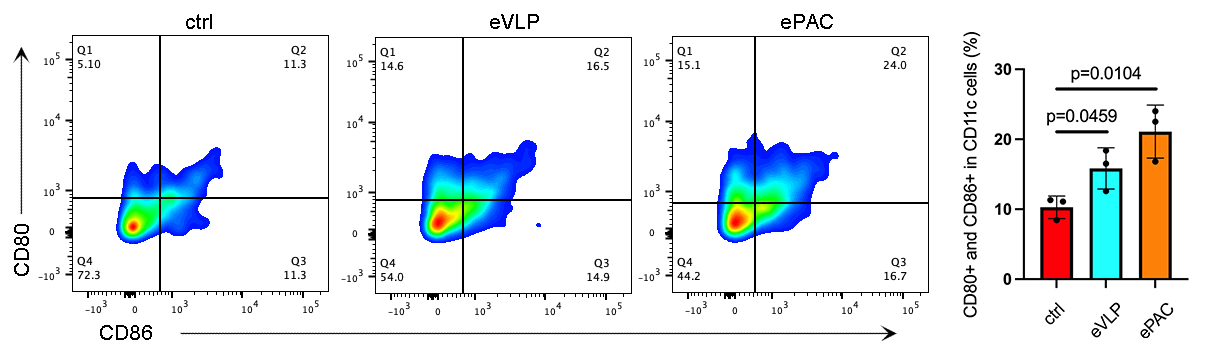

**Supplementary Figure 4. The co-expression of CD80 and CD86 in DCs treated by different formulations and detected by flow cytometry.**

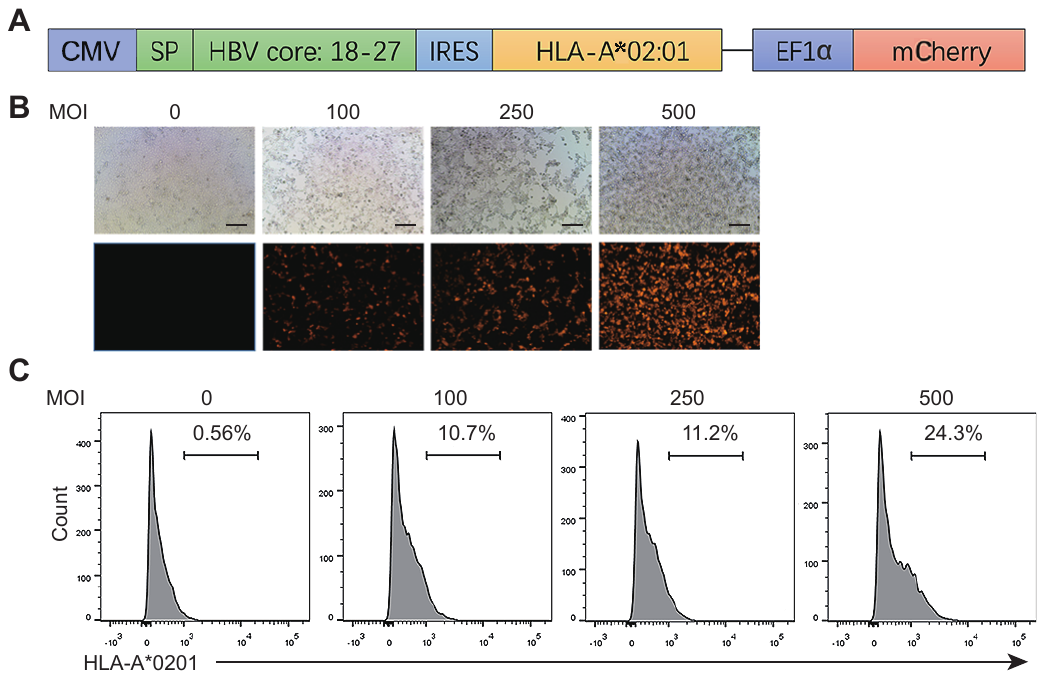

**Supplementary Figure 5. Assessment of the expression of HLA-A*0201 in Hepa1-6 cells**

(A) The plasmid structure of pDC315 for adenovirus package. SP: signal peptide from HLA. (B) The fluorescent images from Hepa1-6 cells infected by adenovirus at various Multiplicity of Infection (MOI). (C) The expression of HLA-A*0201 in Hepa1-6 cells from (B).

**Supplementary Table 1. The sequences of plasmids used in the study**

| 1. pcDNA3.1-gag-//-eGFP-HA: the sequence of gag-//-eGFP-HA as shown in right was inserted into the pCDH back bone through BglII and XhoI. | agatctgcattcgccaccatggctgctgcaggtggttcatcaaactgcccgccccctccccctccccctcctcccaacaacaacaacaacaacaacaccccaaagagcccaggcgtgcctgacgccgaagatgatgatgaacgcagacacgatgagctccctgaagacatcaacaactttgacgaagacatgaacaggcagtttgagaatatgaacctgctggatcaggtggagttgcttgcacagagctacagtctgctggatcatttagatgactttgatgatgatgatgaagacgatgactttgatccagaacctgaccaggatgagctccctgagtacagtgacgatgatgacctggagcttcagggtgctgcagcagcccctatcccaaactttttctccgatgatgactgccttgaagaccttcctgagaagttcgatggcaaccctgacatgctgggtcctttcatgtatcagtgccagctcttcatggaaaagagcaccagagatttctcagttgaccgcatccgtgtgtgcttcgtgacaagcatgctgatcggccgtgccgcccgctgggctactgccaagctgcaaagatgtacttacctgatgcacaactacactgcctttatgatggagctgaagcatgtctttgaagaccctcagagacgtgaagctgccaaacgcaagatcagacgtctgcgccagggccctgggcctgttgtggactactccaatgcattccagatgattgcccaggacctggattggactgagcctgccctgatggatcagttccaggaaggtctcaacccagacattcgcgcagagctgtctcgccaggaggcccccaagaccctggctgctctgattactgcctgtattcacatcgagagaaggctggctcgtgacgctgctgcaaagcccgatccttcacccagagccttggtgatgcctccaaacagccagaccgatcccaccgagcctgtgggaggtgcccgcatgcgcctgtccaaggaagaaaaggagagacgccgcaaaatgaatttgtgtctctactgtggcaatggaggccatttcgccgacacgtgtccagcgaaagcctccaagaattcgatggtgagcaagggcgaggagctgttcaccggggtggtgcccatcctggtcgagctggacggcgacgtaaacggccacaagttcagcgtgtccggcgagggcgagggcgatgccacctacggcaagctgaccctgaagttcatctgcaccaccggcaagctgcccgtgccctggcccaccctcgtgaccaccctgacctacggcgtgcagtgcttcagccgctaccccgaccacatgaagcagcacgacttcttcaagtccgccatgcccgaaggctacgtccaggagcgcaccatcttcttcaaggacgacggcaactacaagacccgcgccgaggtgaagttcgagggcgacaccctggtgaaccgcatcgagctgaagggcatcgacttcaaggaggacggcaacatcctggggcacaagctggagtacaactacaacagccacaacgtctatatcatggccgacaagcagaagaacggcatcaaggtgaacttcaagatccgccacaacatcgaggacggcagcgtgcagctcgccgaccactaccagcagaacacccccatcggcgacggccccgtgctgctgcccgacaaccactacctgagcacccagtccgccctgagcaaagaccccaacgagaagcgcgatcacatggtcctgctggagttcgtgaccgccgccgggatcactctcggcatggacgagctgtacaagtatccgtatgatgttccggattatgcatagtaactcgag |
| --- | --- |
| 2. pcDNA3.1-gag-GS-eGFP-HA: the sequence of gag-GS-eGFP-HA as shown in right was inserted into the pCDH back bone through BglII and XhoI. | agatctgcattcgccaccatggctgctgcaggtggttcatcaaactgcccgccccctccccctccccctcctcccaacaacaacaacaacaacaacaccccaaagagcccaggcgtgcctgacgccgaagatgatgatgaacgcagacacgatgagctccctgaagacatcaacaactttgacgaagacatgaacaggcagtttgagaatatgaacctgctggatcaggtggagttgcttgcacagagctacagtctgctggatcatttagatgactttgatgatgatgatgaagacgatgactttgatccagaacctgaccaggatgagctccctgagtacagtgacgatgatgacctggagcttcagggtgctgcagcagcccctatcccaaactttttctccgatgatgactgccttgaagaccttcctgagaagttcgatggcaaccctgacatgctgggtcctttcatgtatcagtgccagctcttcatggaaaagagcaccagagatttctcagttgaccgcatccgtgtgtgcttcgtgacaagcatgctgatcggccgtgccgcccgctgggctactgccaagctgcaaagatgtacttacctgatgcacaactacactgcctttatgatggagctgaagcatgtctttgaagaccctcagagacgtgaagctgccaaacgcaagatcagacgtctgcgccagggccctgggcctgttgtggactactccaatgcattccagatgattgcccaggacctggattggactgagcctgccctgatggatcagttccaggaaggtctcaacccagacattcgcgcagagctgtctcgccaggaggcccccaagaccctggctgctctgattactgcctgtattcacatcgagagaaggctggctcgtgacgctgctgcaaagcccgatccttcacccagagccttggtgatgcctccaaacaagaattcgggaggcggagggagcggaggcggagggagtggaggcggcggatctatggtgagcaagggcgaggagctgttcaccggggtggtgcccatcctggtcgagctggacggcgacgtaaacggccacaagttcagcgtgtccggcgagggcgagggcgatgccacctacggcaagctgaccctgaagttcatctgcaccaccggcaagctgcccgtgccctggcccaccctcgtgaccaccctgacctacggcgtgcagtgcttcagccgctaccccgaccacatgaagcagcacgacttcttcaagtccgccatgcccgaaggctacgtccaggagcgcaccatcttcttcaaggacgacggcaactacaagacccgcgccgaggtgaagttcgagggcgacaccctggtgaaccgcatcgagctgaagggcatcgacttcaaggaggacggcaacatcctggggcacaagctggagtacaactacaacagccacaacgtctatatcatggccgacaagcagaagaacggcatcaaggtgaacttcaagatccgccacaacatcgaggacggcagcgtgcagctcgccgaccactaccagcagaacacccccatcggcgacggccccgtgctgctgcccgacaaccactacctgagcacccagtccgccctgagcaaagaccccaacgagaagcgcgatcacatggtcctgctggagttcgtgaccgccgccgggatcactctcggcatggacgagctgtacaagtatccgtatgatgttccggattatgcatagtaactcgag |
| 3. pcDNA3.1-gag-//-pol-eGFP-HA: the sequence of gag-//-pol-eGFP-HA as shown in right was inserted into the pCDH back bone through BglII and XhaI. | agatctgcattcgccaccatggctgctgcaggtggttcatcaaactgcccgccccctccccctccccctcctcccaacaacaacaacaacaacaacaccccaaagagcccaggcgtgcctgacgccgaagatgatgatgaacgcagacacgatgagctccctgaagacatcaacaactttgacgaagacatgaacaggcagtttgagaatatgaacctgctggatcaggtggagttgcttgcacagagctacagtctgctggatcatttagatgactttgatgatgatgatgaagacgatgactttgatccagaacctgaccaggatgagctccctgagtacagtgacgatgatgacctggagcttcagggtgctgcagcagcccctatcccaaactttttctccgatgatgactgccttgaagaccttcctgagaagttcgatggcaaccctgacatgctgggtcctttcatgtatcagtgccagctcttcatggaaaagagcaccagagatttctcagttgaccgcatccgtgtgtgcttcgtgacaagcatgctgatcggccgtgccgcccgctgggctactgccaagctgcaaagatgtacttacctgatgcacaactacactgcctttatgatggagctgaagcatgtctttgaagaccctcagagacgtgaagctgccaaacgcaagatcagacgtctgcgccagggccctgggcctgttgtggactactccaatgcattccagatgattgcccaggacctggattggactgagcctgccctgatggatcagttccaggaaggtctcaacccagacattcgcgcagagctgtctcgccaggaggcccccaagaccctggctgctctgattactgcctgtattcacatcgagagaaggctggctcgtgacgctgctgcaaagcccgatccttcacccagagccttggtgatgcctccaaacagccagaccgatcccaccgagcctgtgggaggtgcccgcatgcgcctgtccaaggaagaaaaggagagacgccgcaaaatgaatttgtgtctctactgtggcaatggaggccatttcgccgacacgtgtccagcgaaagcctccaagaattcgccgccgggaaactccccggccccgctgtagggggaccttcagcgacagggccagaacgaataaggtccccaccctccgaggcttcgactcagcacctgcaagtgatgctccagattcatatgccgggcagacccaccctgtttgtccgagctatgattgattctggtgcatctggcaacttcattgatcaagactttgtcatacaaaatgcaattcctctcagaatcaaagactggccagtgatggtggaagctattgatgggcatccaattgcctcgggcccaatcattttggaaacccaccacctgatagttgatctgggagaccaccgtgagatactgtcatttgatgtgactcagtctccattctttcctattgtcctaggaatAcgttggctctctactcacgacccacacatcacctggagtactcgctccatagtcttcaactctgattactgtcggttgagatgccgaatgttcgctcagatcccaagcaacctgctctttaccgttccccaaccaaatctccacccctatctcctgcatcatgtacacccccatgtacaccctcacatgcatcaacaccttcatcaacacctgcatcagtttcttcatcctgatccacatcagtatccacacccagacccccattaccatcatcaccagcaggctgatatgcagcatcagctgcaacagtatttgtatcagtacttgtactaccatctgtaccctgtcatgcatcatcatcttcctcctgaccaacatgaacatctgcacgaatatcttcaccaatacctccatcagtaccttcaccaattcctccaccaccatcttcaccctgacttgcaccaatatttgtaccagtatcttcacaaccacatgaatccagatccacatcaccatccccatccagatccccctcaggatccacatcaccctccacatcaggatccccatcaggatcctccacatcaggatccacatcaggatgcacatcaggatccccatatggatccacacctgcatcagcaccagcatccgcagccgcagccgcatccacaacagcatcctaaccatcctcagcagccaccattcttctaccacatggctggattcagaatttaccaccctgtaaggtattactatattcagaatgtgtatacacctgttgatgagcatgtctatccgggtcaccgggtggttgaccctaacattgagatgattcctggagcgcacagcctgcccagtggacatttgtactcaatgtctgagtctgaaatgaatgctctgcgaaatttcgtggacaggaatgttaaagatgggctcatgactcccactgtggcgcccaatggagcccaagtcctgcaagtgaaaagagggtggaaactccaagtcacttacaattgccgagctccacagagtggcaccatccaaaatcagtacctacgcatgtctcttccaaatatgggagaccctgcacacctggcaagctatggtgaatttgtccaagttcctggctacccatatccagcctatgtttactatacaagcccgcatatgatgactgcgtggtacccagtaggacgagatgtacatggacgaataatcgttgtgcctgttgtaatcacctggtctcaaaatacgaaccgccagcctccggtgccccagtatcctcctccgcagccacctccaccaccaccaccacctccaccgccaccaccacctccaccagcatcatcctgcagtgctgcgggaggcggagggagcggaggcggagggagtggaggcggcggatctctcgaggccaccatggtgagcaagggcgaggagctgttcaccggggtggtgcccatcctggtcgagctggacggcgacgtaaacggccacaagttcagcgtgtccggcgagggcgagggcgatgccacctacggcaagctgaccctgaagttcatctgcaccaccggcaagctgcccgtgccctggcccaccctcgtgaccaccctgacctacggcgtgcagtgcttcagccgctaccccgaccacatgaagcagcacgacttcttcaagtccgccatgcccgaaggctacgtccaggagcgcaccatcttcttcaaggacgacggcaactacaagacccgcgccgaggtgaagttcgagggcgacaccctggtgaaccgcatcgagctgaagggcatcgacttcaaggaggacggcaacatcctggggcacaagctggagtacaactacaacagccacaacgtctatatcatggccgacaagcagaagaacggcatcaaggtgaacttcaagatccgccacaacatcgaggacggcagcgtgcagctcgccgaccactaccagcagaacacccccatcggcgacggccccgtgctgctgcccgacaaccactacctgagcacccagtccgccctgagcaaagaccccaacgagaagcgcgatcacatggtcctgctggagttcgtgaccgccgccgggatcactctcggcatggacgagctgtacaagtatccgtatgatgttccggattatgcatagtaatctaga |
| 4. pcDNA3.1-gag-neoantigens: the sequence of gag-neoantigens as shown in right was inserted into the pCDH back bone through BglII and XhoI. | agatctgcattcgccaccatggctgctgcaggtggttcatcaaactgcccgccccctccccctccccctcctcccaacaacaacaacaacaacaacaccccaaagagcccaggcgtgcctgacgccgaagatgatgatgaacgcagacacgatgagctccctgaagacatcaacaactttgacgaagacatgaacaggcagtttgagaatatgaacctgctggatcaggtggagttgcttgcacagagctacagtctgctggatcatttagatgactttgatgatgatgatgaagacgatgactttgatccagaacctgaccaggatgagctccctgagtacagtgacgatgatgacctggagcttcagggtgctgcagcagcccctatcccaaactttttctccgatgatgactgccttgaagaccttcctgagaagttcgatggcaaccctgacatgctgggtcctttcatgtatcagtgccagctcttcatggaaaagagcaccagagatttctcagttgaccgcatccgtgtgtgcttcgtgacaagcatgctgatcggccgtgccgcccgctgggctactgccaagctgcaaagatgtacttacctgatgcacaactacactgcctttatgatggagctgaagcatgtctttgaagaccctcagagacgtgaagctgccaaacgcaagatcagacgtctgcgccagggccctgggcctgttgtggactactccaatgcattccagatgattgcccaggacctggattggactgagcctgccctgatggatcagttccaggaaggtctcaacccagacattcgcgcagagctgtctcgccaggaggcccccaagaccctggctgctctgattactgcctgtattcacatcgagagaaggctggctcgtgacgctgctgcaaagcccgatccttcacccagagccttggtgatgcctccaaacagccagaccgatcccaccgagcctgtgggaggtgcccgcatgcgcctgtccaaggaagaaaaggagagacgccgcaaaatgaatttgtgtctctactgtggcaatggaggccatttcgccgacacgtgtccagcgaaagcctccaagaattcgggaggcggagggagcggaggcggagggagtggaggcggcggatctatgaaggcccgaaactacctgcagtttctgccctcgaaaaccaaggtggctggaggcggagggagccgaggagaacactaccggtacaaggtcagcctccccgggggccagcacgccggaggcggagggagccatgttctctgggacttaaagcagatgtttcggtgtgctgtcttgaaaaacggaggcggagggagctgggacacttgtaccacttacaagtggcaaaagacactggaaggtcatgatggaggcggagggagcctatcaacgtacagaacagcttgcacgttacgatttgtacagaagcgatgcggaggcggagggagcctgtacactcacttcctgcagttgccactggcagccaccgggttctccgtgggaggcggagggagcaaaaggtggttatattggcaacctactctcactaagatggggtttgtgtcatgataactcgag |
| 5. pCDH-DEC-205-mCherry: the sequence of DEC-205-mCherry as shown in right was inserted into the pCDH back bone through XbaI and NotI. | tctagagccaccatgcggacgggccgggtgaccccgggcctggcggcggggctactcctgctgttgctgcggtccttcgggcttgtggagccttctgagagctcaggtaatgatccattcaccatcgtccatgaaaacactggcaagtgcatccagccgctgtctgactgggtagtggcccaggactgtagcggaactaacaacatgttgtggaagtgggtgtcccagcaccgcctctttcacctggaatcccagaagtgcctcggcctcgatattaccaaagccacggacaacctgcgaatgttcagctgtgactccaccgtcatgctgtggtggaaatgtgagcaccattcgctgtacaccgctgcccagtacaggctagctctgaaagatggatatgccgtagccaatacgaatacatctgatgtctggaagaagggaggctccgaggaaaacctttgtgcccagccttatcatgagatatacaccagagatgggaattcctacgggagaccttgtgaattccctttcttgattggtgagacatggtaccatgactgcattcatgatgaagatcatagtgggccatggtgtgccactaccctaagttatgaatatgatcaaaagtggggcatctgcctactaccagaaagtggctgtgaaggtaactgggaaaagaatgagcagattggaagttgctaccaatttaataatcaggaaattctgtcttggaaagaagcttatgtttcctgtcagaaccaaggagctgacttactgagcatccacagtgctgccgaattagcctacattacgggaaaagaggacattgctagacttgtttggcttggactgaatcagctctattctgcgagaggttgggaatggtcagacttcaggccactcaaatttcttaactgggatccaggcacgcccgttgcacctgtgattggtgggtcaagctgtgccagaatggacacagagtccgggctgtggcaaagtgtttcctgtgaatctcagcagccttacgtctgcaagaagccactgaacaacacgctggagctcccagatgtttggacttacacagatacccactgccatgtgggctggctgccaaataatgggttttgctatctgctggcgaatgaaagtagttcctgggatgcagcacatttgaaatgcaaagccttcggtgcagacctcatcagcatgcactccttagcagatgtggaggtggttgtcacgaaactccataatggggatgtcaaaaaagaaatatggacaggccttaaaaacacaaacagccctgctttgttccagtggtcggacggaacggaagttactctaacgtactggaatgagaatgagccgagtgttcccttcaacaagactcccaactgtgtttcctatttaggaaagttaggtcagtggaaagtccagtcctgtgagaagaaactcagatatgtatgcaagaaaaagggagaaataactaaggatgcagagtcggataagctgtgtccgccagacgagggctggaagagacatggagaaacctgttacaagatttatgagaaagaggcccctttcggaacgaactgcaacctgaccatcactagcaggttcgagcaggaattcttgaattatatgatgaagaactatgataagtcccttcggaagtacttctggactggcctgagagaccctgactctcgaggagaatacagttgggccgttgctcagggagtaaagcaggctgtgaccttttccaactggaattttcttgaaccggcgtctccaggcgggtgcgtggctatgtctactggaaagactcttggcaagtgggaagtgaagaactgcagaagcttccgtgctctttcaatatgcaagaaagtgagcgaaccccaggagcctgaagaagcagcccccaagcccgacgacccctgtcctgaaggctggcacactttcccctccagcctttcttgttataaggtgttccatatagaaagaatcgtaagaaagaggaactgggaagaagccgaaaggttctgccaagcccttggagctcacctacccagcttcagtcgtagagaggaaattaaggactttgtgcatttgttaaaggaccagttcagtgggcagcgttggttgtggattggtctgaataagagaagccctgatttacaagggtcctggcagtggagtgaccggacaccagtgtctgctgtgatgatggagccggagtttcaacaggattttgacatcagagactgtgctgccatcaaggtccttgatgtaccttggcgaagagtctggcatctctatgaggacaaggactatgcttactggaaaccttttgcttgtgatgccaagcttgagtgggtgtgccagattccaaaaggtagcactccccagatgccagactggtataatccagagcgcactggaattcatgggcccccagttataattgaaggaagtgaatactggtttgttgctgatccccacttaaactacgaagaagccgtcttatactgtgctagcaatcacagctttcttgccacgataacatcgttcacaggactaaaagctatcaaaaacaaactagcaaatatttctggcgaggaacagaagtggtgggtgaaaacgagtgagaatccaattgatcgttactttctaggctcgcgccgccgcctgtggcaccatttccccatgacgtttggagatgaatgtttgcacatgtcagccaagacgtggcttgttgacttaagtaaacgagcggactgtaatgccaagttgcccttcatctgtgaaagatacaatgtctcttcattagagaaatacagcccagatcctgcagccaaagtacagtgcactgagaagtggattccttttcaaaataagtgcttcctaaaggtcaactctgggcccgttacgttttctcaagcaagcggcatttgtcattcctacggcggcacccttccttccgtgctgagccggggtgaacaagatttcattatatccttgcttcctgaaatggaagctagtctatggattggtctgcgctggactgcctacgaaaggataaacagatggacagacaacagagagctgacctacagcaactttcacccactgctggtcggtcggaggctgagcataccaacgaatttctttgatgatgagtcccacttccactgcgccttgattcttaatctcaaaaagtcaccgcttactgggacctggaattttacttcctgttcagaacgacactctctgtctctctgtcaaaaatactcagagactgaagacggacagccctgggagaacacttcaaaaacagtgaagtatctaaataacctatacaaaatcatctcgaagcccctgacgtggcacggcgctctgaaggagtgcatgaaagagaagatgaggttggtgagcatcacagacccttaccagcaggccttcctcgcagtgcaggccaccctgcgcaacagctccttctggatcggactctccagtcaagatgatgaactcaactttggttggtcagatgggaaacgtcttcaatttagtaactgggctggaagcaatgagcaacttgatgactgcgtgatattagacacagatggattctggaaaacagctgactgtgatgataaccagcctggcgccatttgctactatccaggaaatgagactgaggaggaggtcagagcactggacactgctaaatgcccgtctcctgtacagagcaccccatggataccattccagaactcctgctacaatttcatgattaccaacaacaggcataagacagtcacaccggaggaagtgcagtccacgtgcgagaagctgcattcgaaagcacacagtctgagcattcggaatgaggaggagaatacctttgttgtggaacagcttctgtacttcaattatattgcctcatgggtcatgttaggaataacctatgaaaacaattctttgatgtggtttgataaaactgcattgtcctacacacactggagaacgggaagaccaactgtgaaaaatggcaaatttttggctggtctaagtactgatggattctgggatattcagtctttcaatgttattgaagaaacacttcatttttaccagcacagtatttctgcttgtaaaattgaaatggttgactatgaggacaaacacaatggcaccctgccacagttcattccatataaggacggcgtctacagcgttattcagaagaaggtgacgtggtatgaagcattgaacgcgtgctctcaaagtgggggagagttggccagtgttcacaacccaaatgggaagctctttctggaagacattgtgaaccgtgacggattccctctctgggttgggctctcaagtcatgatggaagcgaatcgagtttcgaatggtccgatggcagagcatttgactatgtcccatggcagagcctacaatctcccggagactgtgtcgtcttatatccaaaaggaatttggagacgtgaaaaatgcctgtctgttaaggatggtgctatttgttacaagcctacaaaagataaaaagctgatctttcatgtaaaatcatcaaaatgtccagtggcaaagagggatggtccccagtgggtccagtatgggggccactgttacgcttcggaccaggtactgcacagcttctcagaggccaaacaagtgtgtcaagagcttgatcattcggcaactgttgtcaccatagcagatgaaaatgagaataagtttgtgagcagactgatgagggagaactataatattactatgagagtttggcttggcctgtctcagcattcactcgatcagtcttggagttggctcgatggattagatgtgacatttgtcaaatgggaaaataaaactaaggatggtgatgggaaatgtagcattttaatagcttcaaatgaaacctggagaaaagtccattgctcacgtggctatgcaagagctgtctgcaaaattcctctgagcccggactacacaggcatagccatcctgtttgccgtgctgtgcctcttagggctcatcagcttggcgatttggttcctcttgcaacgatcccatatccgctggaccggcttctcctcggttcggtatgaacatggaaccaacgaagacgaggtgatgctcccttctttccacgacaccggtggttctggaggttcaatggtgagcaagggcgaggaggataacatggccatcatcaaggagttcatgcgcttcaaggtgcacatggagggctccgtgaacggccacgagttcgagatcgagggcgagggcgagggccgcccctacgagggcacccagaccgccaagctgaaggtgaccaagggtggccccctgcccttcgcctgggacatcctgtcccctcagttcatgtacggctccaaggcctacgtgaagcaccccgccgacatccccgactacttgaagctgtccttccccgagggcttcaagtgggagcgcgtgatgaacttcgaggacggcggcgtggtgaccgtgacccaggactcctccctgcaggacggcgagttcatctacaaggtgaagctgcgcggcaccaacttcccctccgacggccccgtaatgcagaagaagaccatgggctgggaggcctcctccgagcggatgtaccccgaggacggcgccctgaagggcgagatcaagcagaggctgaagctgaaggacggcggccactacgacgctgaggtcaagaccacctacaaggccaagaagcccgtgcagctgcccggcgcctacaacgtcaacatcaagttggacatcacctcccacaacgaggactacaccatcgtggaacagtacgaacgcgccgagggccgccactccaccggcggcatggacgagctgtacaagtaatgagcggccgc |
| 6. pcDNA3.1-gag-HBc 18-27: the sequence of gag-HBc 18-27 as shown in right was inserted into the pCDH back bone through BglII and XhoI. | agatctgcattcgccaccatggctgctgcaggtggttcatcaaactgcccgccccctccccctccccctcctcccaacaacaacaacaacaacaacaccccaaagagcccaggcgtgcctgacgccgaagatgatgatgaacgcagacacgatgagctccctgaagacatcaacaactttgacgaagacatgaacaggcagtttgagaatatgaacctgctggatcaggtggagttgcttgcacagagctacagtctgctggatcatttagatgactttgatgatgatgatgaagacgatgactttgatccagaacctgaccaggatgagctccctgagtacagtgacgatgatgacctggagcttcagggtgctgcagcagcccctatcccaaactttttctccgatgatgactgccttgaagaccttcctgagaagttcgatggcaaccctgacatgctgggtcctttcatgtatcagtgccagctcttcatggaaaagagcaccagagatttctcagttgaccgcatccgtgtgtgcttcgtgacaagcatgctgatcggccgtgccgcccgctgggctactgccaagctgcaaagatgtacttacctgatgcacaactacactgcctttatgatggagctgaagcatgtctttgaagaccctcagagacgtgaagctgccaaacgcaagatcagacgtctgcgccagggccctgggcctgttgtggactactccaatgcattccagatgattgcccaggacctggattggactgagcctgccctgatggatcagttccaggaaggtctcaacccagacattcgcgcagagctgtctcgccaggaggcccccaagaccctggctgctctgattactgcctgtattcacatcgagagaaggctggctcgtgacgctgctgcaaagcccgatccttcacccagagccttggtgatgcctccaaacagccagaccgatcccaccgagcctgtgggaggtgcccgcatgcgcctgtccaaggaagaaaaggagagacgccgcaaaatgaatttgtgtctctactgtggcaatggaggccatttcgccgacacgtgtccagcgaaagcctccaagaattcgggaggcggagggagctttcttccaagcgacttctttccaagtgtgtgataactcgag |

**Supplementary Table 2. Antibodies used in the study**

| Target | Clone | Source | Usage |
| --- | --- | --- | --- |
| CD11c-APC | N418 | Biolegend (#117310) | Flow cytometry |
| CD80-PE | 16-10A1 | Biolegend (#104708) | Flow cytometry |
| CD86-PeCy7 | GL-1 | Biolegend (#105014) | Flow cytometry |
| MHC-II-PE | M5/114.15.2 | eBioscience (#17-5321-82) | Flow cytometry |
| CD3-FITC | 17A2 | Biolegend (#100204) | Flow cytometry |
| CD8-APC | 53-5.8 | Biolegend (#140410) | Flow cytometry |
| CD44-PeCy7 | IM7 | Biolegend (#103030) | Flow cytometry |
| CD62L-PerCP | MEL-14 | Biolegend (#104430) | Flow cytometry |
| 4-1BB-APC | 17B5 | Biolegend (#106110) | Flow cytometry |
| 4-1BB-APC | 4B4-1 | BD Pharmingen (#550890) | Flow cytometry |
| HLA-A*02-FITC | BB7.2 | Abcam (#ab27728) | Flow cytometry |
| CD4 | EPR19514 | Abcam (#ab183685) | Immunohistochemistry |
| CD8 | EPR21769 | Abcam (#ab217344) | Immunohistochemistry |
| CD8 | EPR21769 | Abcam (#ab217344) | Immunofluorescence |
| PD-1 | EPR20665 | Abcam (#ab214421) | Immunofluorescence |
| TIM-3 | EPR22241 | Abcam (#ab241332) | Immunofluorescence |
| CTLA-4 | CAL49 | Abcam (#ab237712) | Immunofluorescence |
| HA tag | C29F4 | CST (#3724) | Western Blot |
| GFP | D5.1 | CST (#2956) | Western Blot |
| P65 | D14E12 | CST (#8242) | Western Blot |
| phospho-P65 | 93H1 | CST (#3033) | Western Blot |
| Myd88 | D80F5 | CST (#4283) | Western Blot |
| Actin | 13E5 | CST (#4970) | Western Blot |
| CD9 | E8L5J | CST (#98327) | Western Blot |
